# Re-discovery of PF-3845 as a new chemical scaffold inhibiting phenylalanyl-tRNA synthetase in *Mycobacterium tuberculosis*

**DOI:** 10.1101/2020.10.27.357681

**Authors:** Heng Wang, Min Xu, Curtis A Engelhart, Xi Zhang, Baohua Yan, Miaomiao Pan, Yuanyuan Xu, Shilong Fan, Renhe Liu, Lan Xu, Lan Hua, Dirk Schnappinger, Shawn Chen

**Author notes:** Corresponding author: Shawn Chen. These authors contributed equally to this work (Heng Wang, Min Xu).

## Abstract

*Mycobacteria tuberculosis* (Mtb) remains the deadliest pathogenic bacteria worldwide. The search for new antibiotics to treat drug-sensitive as well as drug-resistant tuberculosis has become a priority. The essential enzyme phenylalanyl-tRNA synthetase (PheRS) is an antibacterial drug target because of the large differences between bacterial and human PheRS counterparts. In a high-throughput screening of 2148 bioactive compounds, PF-3845, which is a known inhibitor of human fatty acid amide hydrolase (FAAH), was identified inhibiting Mtb PheRS at *K*_*i*_ ∼0.73 ± 0.055 µM. The inhibition mechanism was studied with enzyme kinetics, protein structural modelling and crystallography, in comparison to a PheRS inhibitor of the noted phenyl-thiazolylurea-sulfonamide class. The 2.3-Å crystal structure of Mtb PheRS in complex with PF-3845 revealed its novel binding mode, in which a trifluoromethyl-pyridinylphenyl group occupies the Phe pocket while a piperidine-piperazine urea group binds into the ATP pocket through an interaction network enforced by a sulfate ion. It represents the first non-nucleoside bi-substrate competitive inhibitor of bacterial PheRS. PF-3845 inhibits the *in vitro* growth of Mtb H37Rv at ∼24 µM, and the potency of PF-3845 increased against Mtb pheS-FDAS, suggesting on target activity in mycobacterial whole cells. PF-3845 does not inhibit human cytoplasmic or mitochondrial PheRSs in biochemical assay, which can be explained from the crystal structures. Further elaboration of the piperidine-piperazine urea moiety by medicinal chemistry effort will produce potential antibacterial lead with improved selectivity on the cellular level.

## Introduction

Protein synthesis is the cellular process targeted by many commercial antibiotics. It has always been a focal point of modern antibacterial drug discovery (1). Aminoacyl-tRNA synthetases (aaRSs), a family of ∼20 essential enzymes, ligate amino acids to the corresponding tRNAs that decode messenger RNA to produce protein at the translating macromolecular ribosome (2). Inhibition of bacterial aaRS blocks the translation and ultimately shuts down protein synthesis, which is crucial for pathogens to survive in host or inside host cells (3, 4). Between a bacterial aaRS protein and its human counterpart, either the large sequence difference or small variation of key residues in the catalytic core explains the high selectivity of successful and promising aaRS inhibitor drugs, as exemplified by the mupirocin used for the treatment of staphylococcal infection (4). Tuberculosis (TB) caused by the single agent *Mycobacterium tuberculosis* (Mtb) has surpassed AIDS/HIV, becoming a leading infectious disease worldwide (5). Most TB drugs were discovered in the past century and are losing efficacy due to the resistance inevitably arisen in bacteria. New chemical scaffold with novel inhibition mechanism against Mtb is being actively sought. An oxaborole compound GSK3036656 that inhibits Mtb leucyl-tRNA synthetase is currently undergoing clinical trial (6).

Bacterial phenylalanyl-tRNA synthetase (PheRS) is a member of class II aaRS based on the active site topology (7). A functional PheRS is typically made of two heterodimers (αβ)_2_, with a whole molecular weight around 250 kD. In Mtb H37Rv genome, two consecutive essential genes *Rv1649* and *Rv1650* (*pheST*) encode the protein subunits (8). Like other aaRSs, PheRS catalyses the formation of phenylalanyl-tRNA^Phe^ in two steps: 1) activation of phenylalanine by hydrolysing ATP to form Phe-AMP and pyrophosphate (PPi); 2) subsequent transfer of Phe to the 2’-OH group of adenosine ribose at the 3’-terminal of tRNA^Phe^, and simultaneous release of AMP. In addition to the synthetic site on the α subunit, PheRS has a dedicated domain on the β subunit that can hydrolyze improperly charged tRNA, e.g. tyrosinyl-tRNA^Phe^, to maintain the fidelity of aminoacylation and translation (9). From a drug discovery standing point, PheRS synthetic activity that involves binding of the three substrates, and the editing activity could all be targetable. PheRS should also allow specific binding of compounds that allosterically regulate the enzymatic activities because of its size, molecular mechanism and other miscellaneous functions (10). PheRS was a major antibacterial target in a number of screening programs (11–18). Bacterial PheRS-specific hits such as phenyl-thiazolylurea-sulfonamides have emerged from synthetic compound library. To the best of our knowledge, most of them were not for TB drug except one that identified a fungal natural product isopatulin able to inhibit the growth of Mtb strain H37Rv (19). Recently, PheRS from *Plasmodium falciparum* that is another pathogen of global health concern, has become genetically and chemically validated target for antiparasitics (20). Potent PheRS inhibitors with novel mechanisms at the molecular and cellular levels were discovered (21), prompting us to initiate a hit discovery campaign targeting Mtb PheRS with new assays and diversity compound libraries, such as recent collection of pharmacologically relevant small molecules (22).

In this work, Mtb PheRS enzyme was enzymatically characterized, and analysed with a synthesized reference compound GDI05-001. High throughput screening (HTS) assay was adopted to screen a Selleck-2148 library against the PheRS. One screening hit, PF-3845 was confirmed to inhibit Mtb PheRS with *K*_i_ ∼0.73 µM. It is a well-developed inhibitor of human fatty acid amide hydrolase (FAAH). Its MIC (minimum concentration required to inhibit >90% growth of wild-type H37Rv) is ∼24 µM and the antibiotic activity increased against Mtb pheS-FDAS, in which PheS contained a C-terminal degradation tag. The mechanism of inhibition was studied with enzyme kinetics and crystal structures. PF-3845 was shown to be a non-nucleoside bi-substrate competitive inhibitor of the PheRS. The results will aid future medicinal chemistry effort to improve the potency, reduce the toxicity, and develop combination therapy with TB drugs.

## Results

### Overexpression, purification and characterization of Mtb PheRS

In Mtb H37Rv genome, genes *pheST* encode the α, β subunits of PheRS, respectively. The deduced amino acid sequence of PheS is ∼16%-30% identical to human cytoplasmic hFARS1α or mitochondrial hFARS2. The mitochondrial PheRS does not have a β subunit. The identity of PheT *versus* hFARS1β is below 12%. Three conserved signature motifs of class II aaRS are identified in Mtb PheS sequence (Supporting information Fig S2). The ∼3.5 kb *pheST* was synthesized based on the DNA sequences in GenBank. After sequencing confirmation, the DNA was cloned for heterologous expression in *E. coli*. Several constructs were made for co-purification of the heterodimeric protein. The best one pET30a-MtbPheRS-CHis, under optimized fermentation conditions, gave a yield of ∼1-2 mg/L, with the highest purity possible after optimized purification steps (Supporting information Fig S1B). We also constructed, overexpressed and partially purified a Mtb tRNA_GAA_^Phe^ from *E. coli* (see Experimental Procedures). It was reported more than ∼90% of the enriched and partially purified tRNA fraction could be the overexpressed tRNA species (23, 24). We used standard aminoacylation assay that detects the charging of ^14^C labelled phenylalanine (Phe) onto tRNA^Phe^, to confirm the purified protein and tRNA were indeed functional together. Michaelis-Menten kinetic parameters of Mtb PheRS were measured with the tRNA aminoacylation assay and shown in supporting information Fig S3. The *K*_m_’s and *k*_cat_’s generally agree with previous reports on Mtb’s and other bacterial PheRSs although some discrepancies are noted. For example, we and others found the *K*_m_ with regard to Phe at ∼68 ± 6.6 µM, but this is much higher than the ∼1-7 µM of other bacterial PheRSs (15, 16, 25, 26). The parameters are subject to revision in further research of PheRS biology and aromatic amino acid metabolism.

### Assay development and screening for inhibitors of enzymatic activity

Non-radioactive biochemical assays were adopted for screening for PheRS inhibitor and the subsequent analysis of mode of inhibition. In a continuous spectrophotometric assay (27), the PPi production is coupled to generation of phosphate by inorganic pyrophosphatase; the phosphate then serves as a substrate in a purine nucleoside phosphorylase (PNPase) catalyzed reaction to cleave nucleoside compound MESG into ribose-1-phosphate, and base AMMP that has absorbance at 360 nm. When optimizing the so-called PPi production assay, we noticed the A_360_ readout is tRNA-dependent as expected, meaning no PPi was released in the absence of tRNA (Fig. 1A). PheRS concentration at 100 nM per reaction gave a linear response in the first 6 minutes of the progression. By measuring the initial velocity of PPi production, the apparent *K*_m, ATP_ with regard to ATP was found to be 162.5 ± 11.62 µM, *K*_m, Phe_ 18.1 ± 2.44 µM, and *K*_m, tRNA_ 0.13 ± 0.049 µg/ml (Fig. 1B-D). These *K*_m_’s were used as references in the following screening and hit confirmation.

**FIGURE 1.**
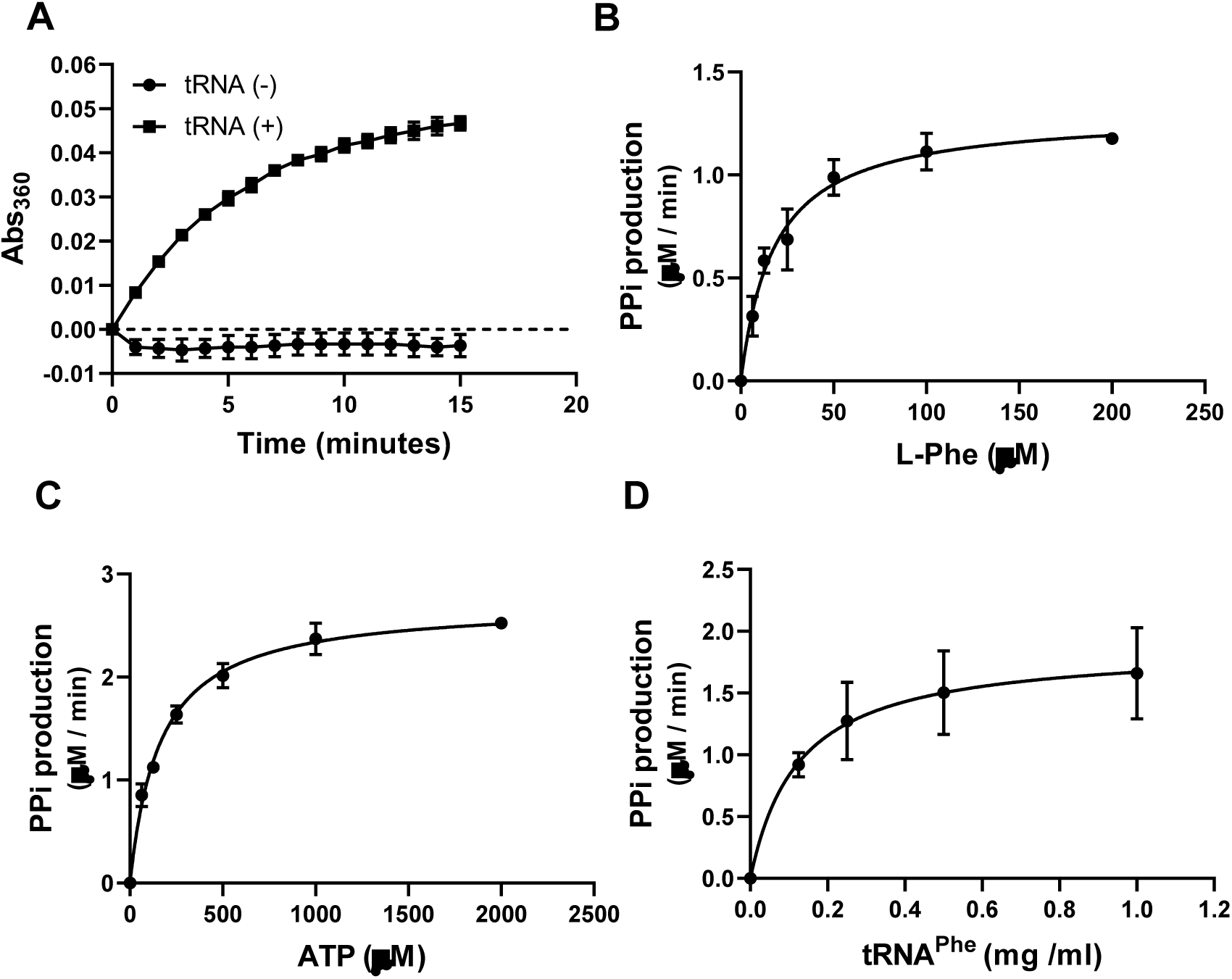
Mtb PheRS pyrophosphate (PPi) production is dependent on tRNA^Phe^ and the Michaelis-Menten kinetics. A, PPi generation in the PheRS reaction with or without tRNA^Phe^. B-D, Km determination of Mtb PheRS with respect to L-Phe, ATP or tRNA^Phe^. All data are presented as mean ± S.D. (n=3).

To screen compound library, we used a commercial kit Kinase-Glo, which measures the remaining amount of ATP in a PheRS reaction. Readout is luminescent signal generated by luciferase catalysed oxidation of luciferin in the presence of the ATP. The readout is thus inversely correlated with the amount of PheRS activity. Because ATP at 1 µM to initiate the PheRS reaction is far below its *K*_*m*_, the assay is presumably biased toward ATP-competitive inhibitors (28). We optimized the assay in 384-well HTS format and determined the amount of PheRS per reaction (Supporting information Fig S4). Reaction endpoint was chosen to be at the first two hours. A known bacterial PheRS inhibitor (12) was synthesized as reference compound, named GDI05-001 (NMR and MS data in supporting information). Its IC_50_ against Mtb PheRS was determined to be 1.724 ± 0. 131 μM.

A Selleck Bioactive Collection purchased before 2018 contains 2148 small molecules with validated biological and pharmacological activities. The library was screened in 7 plates, which showed mean robust Z’ at 0.756-0.862. When the cut-off was set to 70% inhibition compared to no-enzyme control reaction, 4 initial hits were identified (Supporting information Fig. S5), giving a hit rate ∼0.19% (Fig. 2A). A top hit PF-3845 was reported a potent inhibitor of human enzyme FAAH (29) and many analogous compounds are in public database. PF-3845 and one analog PF-04457845 were re-purchased (Fig. 2B). The IC50s of the two compounds were determined with the two assays and other similar assays (30) (Supporting information Fig. S6). By the Kinase-Glo assay, the IC_50_ of PF-3845 was 2.653 ± 0.190 μM (Hillslope −0.860 ± 0.047) and the IC_50_ of PF-04457845 was 9.758 ± 0.726 μM (Hillslope −0.812 ± 0.045) (Fig. 2C). PF-3845, at concentrations up to 20 μM, did not inhibit the activities of pyrophosphatase, PNPase and luciferase that were used in the coupled reactions (Supporting information Fig. S7). The inhibition was observed only when Mtb PheRS was present in the coupled reactions. Thus, we identified investigational drug PF-3845 as a new PheRS inhibitor. PF-3845 apparently has a scaffold different from previously known PheRS inhibitors.

**FIGURE 2.**
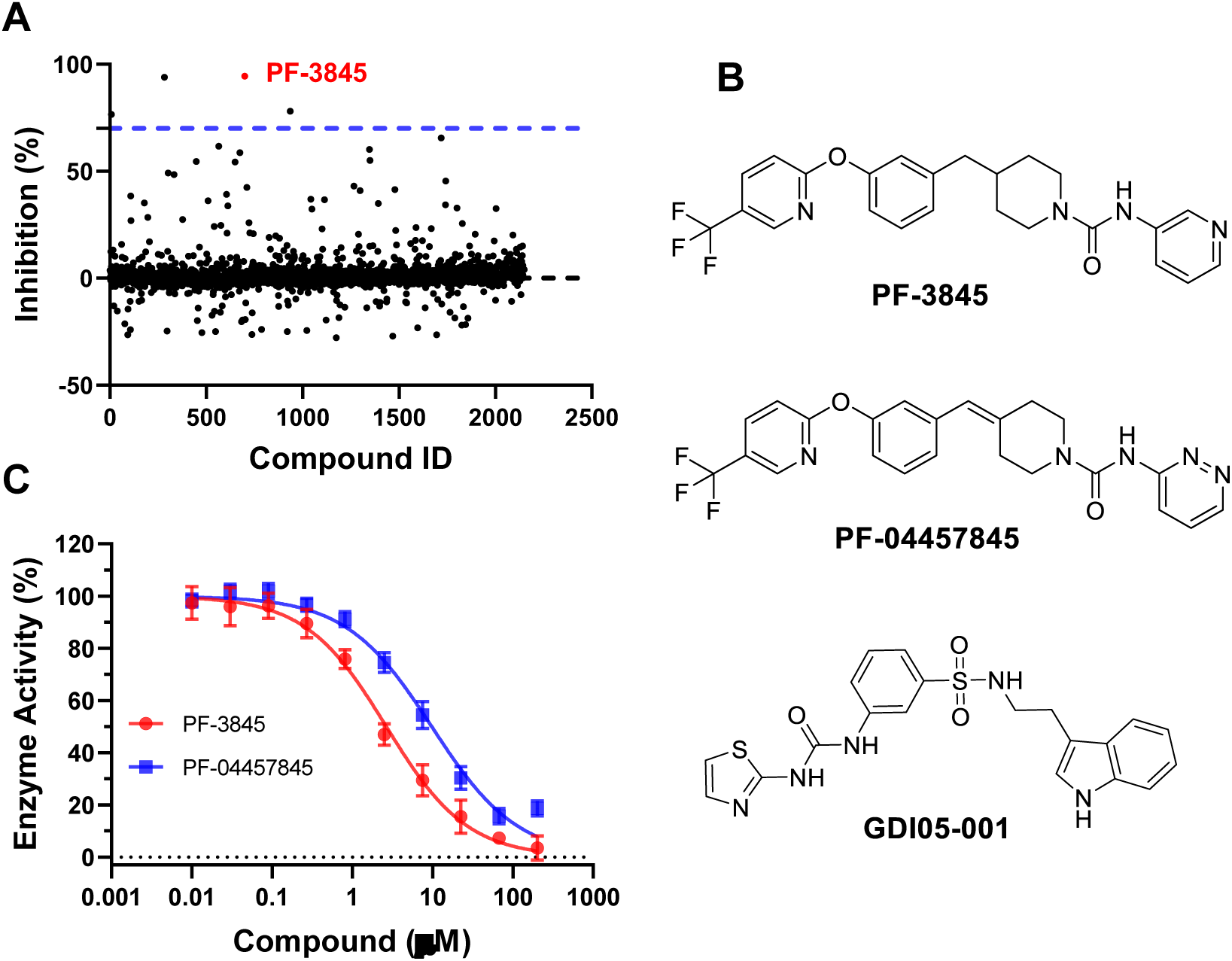
Result of HTS screening with Selleck-2148 library and hit confirmation. A, summary plot of the HTS result. B, structures of PF-3845, a purchased analog PF-04457845 and reference compound GDI05-001. C, IC_50_ of PF-3845 and 004457845 against Mtb PheRS by Kinase-Glo assay, presented as mean ± S.D. (n=3). See supporting information for IC_50_’s determined by other assays.

### Analysis of the mode of inhibition by kinetics, crystallography and modelling

The mode of inhibition of PF-3845 was analyzed with enzyme kinetics in comparison with GDI05-001. PPi production assay was used to record the progress of PheRS reaction setups, which had excessive amounts of two substrates but varying concentration of a third substrate at various inhibitor concentrations. Each dataset was fit into Michaelis-Menten equation; the resulting Lineweaver-Burk plot of a set of experiments with regard to the third substate was examined for a pattern of competitive, non-competitive, uncompetitive or mixed inhibition mode. GDI05-001 and PF-3845 both are clearly phenylalanine competitive inhibitors of PheRS as the presence of inhibitor only affected *K*_*m*_ but not *V*_max_ (Fig. 3A). In the Phe-competitive mode, *K*_*i*_ of GDI05-001 is 0.195 ± 0. 0145 µM and *K*_*i*_ of PF-3845 is 1.70 ± 0.114 µM, determined with the PPi production assay. It was reported GDI05-001 was non-competitive regarding ATP (12) and we had similar observation. In contrast, PF-3845 is more likely in a mixed inhibition mode with regard to ATP over a range of PF-3845 concentrations (Fig. 3B). PF-3845 is an uncompetitive inhibitor with regard to tRNA^Phe^ (Fig. 3C).

**FIGURE 3.**
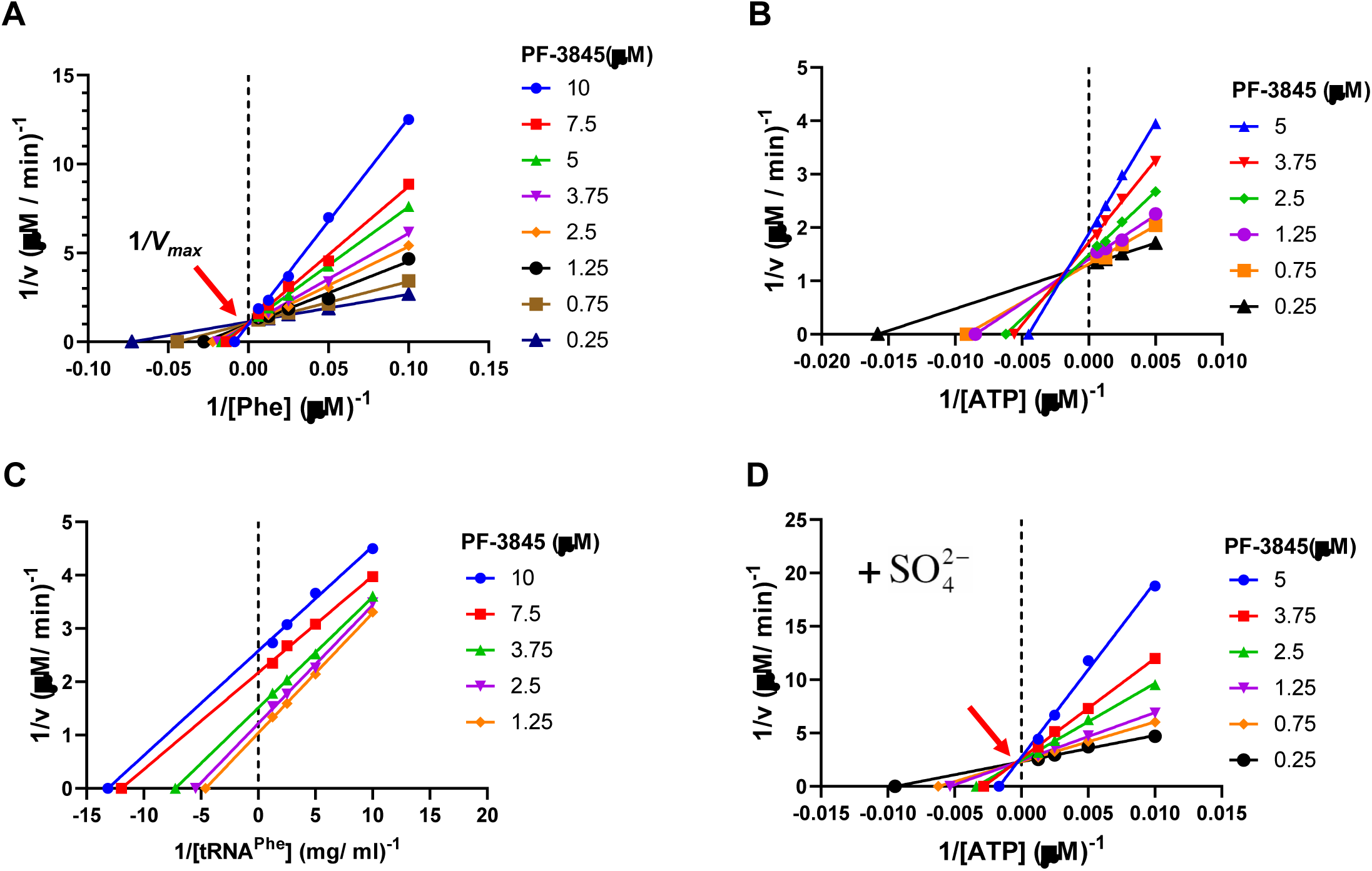
Mode of inhibition of PF-3845, analyzed by Lineweaver-Burke plot with respect to one of the substrates L-Phe (A), ATP (B), tRNA (C) and ATP with SO_4_^2-^ (D) added into the buffer, using the PPi production assay. A, competitive inhibition mode regarding L-Phe. B, mixed inhibition mode regarding ATP. C, uncompetitive inhibition mode regarding tRNA^Phe^. D, competitive inhibition mode regarding ATP, in the presence of 5 mM (NH_4_)_2_SO_4_.

Crystallographic effort was made to first solve the structures of Mtb PheRS apo protein and its complex with GDI05-001 at 2.83 Å and 2.71 Å, respectively. Like other bacterial PheRS structures, Mtb PheRS is a (αβ)_2_ heterotetramer. The α subunit contains catalytic site and β subunit contains editing site (Fig. 4A). Except for the N-terminal 74 residues of α subunit, all other residues were constructed. The N-terminal region of *Thermophilus thermus* PheRS was reported to form a coiled-coil domain, playing a critical role in interaction with tRNA (31). It was flexible and invisible in the solved Mtb PheRS when no tRNA was included. The binding mode of GDI05-001 was illustrated by co-crystallization (Fig. 4B). When looking into the catalytic site of the α subunit, we found that besides the amino acid (Phe) pocket and ATP pocket, there is an additional pocket between the two pockets (Fig. 4C). The key residues forming the additional pocket are mainly hydrophobic, including F143, F148, A154, H175, F254, F255, P256 and F257. GDI05-001 occupies the Phe pocket and the additional pocket while the ATP pocket is empty (Fig. 4C and D). The phenylsulfonamide group of GDI05-001 extends much deeper into the amino acid pocket than a phenylalanine could. In this pocket, strong π-π stackings can be seen between the phenyl rings of GDI05-001 and F255/F257. Three hydrogen bonds are formed between the urea group of GDI05-001 and the side chains of E217 and S177. In addition, the sulfonamide oxygen forms interactions with the side chain of Q215 and the main chain of G282 and G307. The indole moiety of GDI05-001 is bended almost vertically to the phenylsulfonamide and extend into the additional pocket, which has strong hydrophobic interaction with the protein. Compared with the apo form, two major differences were observed. First, in the apo form structure, the side chain of F257 has two alternative conformations with one conformation occupies the deeper amino acid pocket, and the side chain of Q180 also stretches towards the pocket. While in the liganded structure, F257 adapts only one rigid conformation which occupies out of the pocket. Together with the rotation of Q180, they constitute a much deeper amino acid pocket of the complex than the apo form (Supporting information Fig. S8). Second, the loop 148-160 undergoes a conformational change upon GDI05-001 binding and has much higher B factors than the apo structure, indicating it is more flexible when GDI05-001 binds (Supporting information Fig. S9).

**FIGURE 4.**
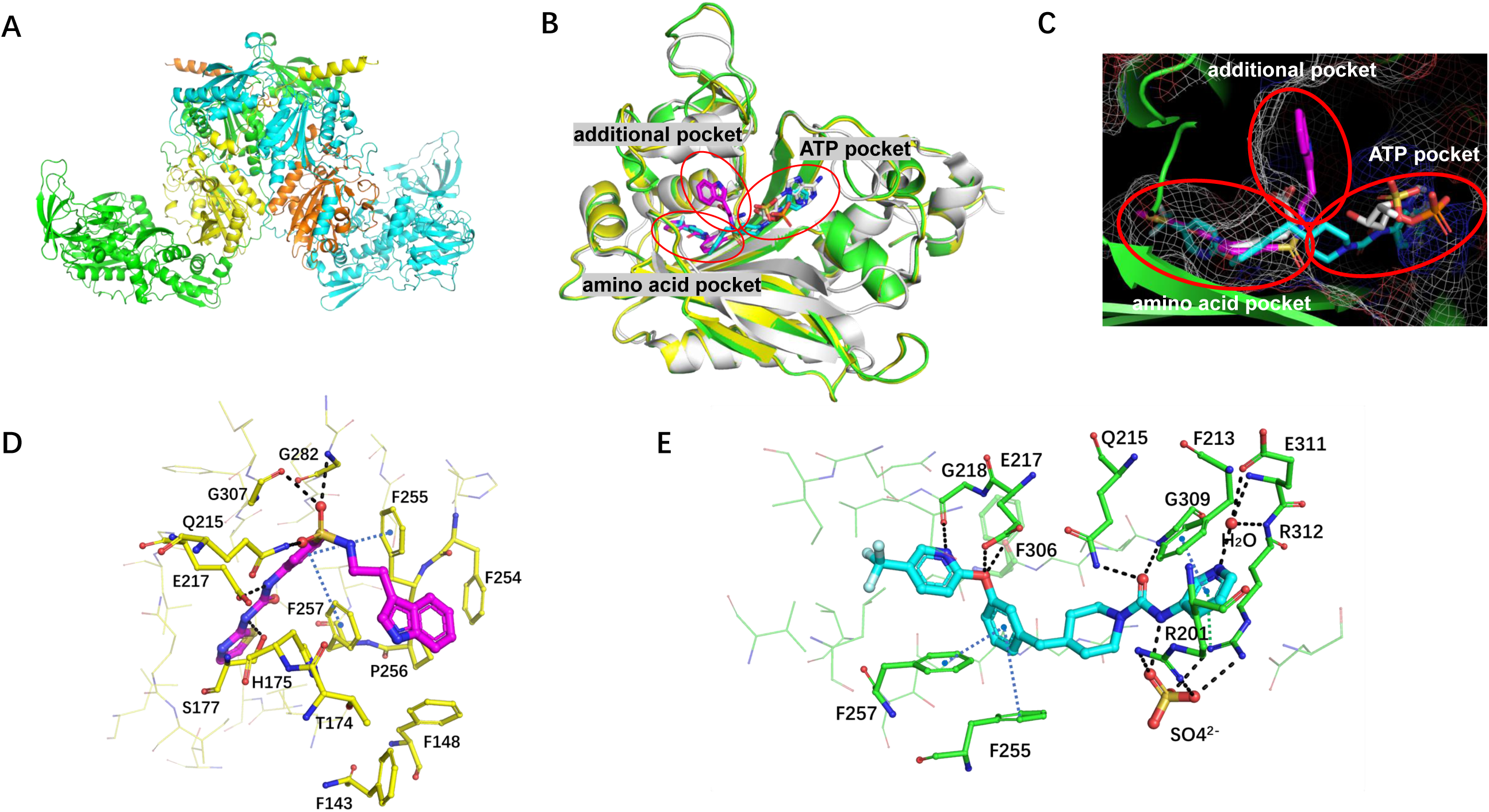
Overall structures of Mtb PheRS, and three binding pockets of the catalytic site. A, overall structure of apo Mtb PheRS. The α subunits are shown in yellow and orange, the β subunits in green and cyan. B, structure superposition of the alpha subunits of MtbPheRS-GDI05-001, MtbPheRS-PF-3845, and EcPheRS-Phe-AMP (PDB: 3PCO). MtbPheRS-GDI05-001 is shown in yellow with ligand in magenta, MtbPheRS-PF-3845 is shown in green with ligand in cyan, EcPheRS-Phe-AMP is shown in white. Three binding pockets of the catalytic site are indicated by circle. C, three binding pockets of the catalytic site are shown in mesh based on the structure of MtbPheRS-PF-3845. D, the interaction network between GDI05-001 and Mtb PheRS. E, the interaction network between PF-3845 and Mtb PheRS. Hydrogen bonds are shown in black dash, π-π stackings are shown in blue dash, π-cation interaction is shown in green dash.

On the other hand, preliminary analysis using a deep learning-based docking platform Orbital showed that PF-3845 might have two major putative binding conformations among the top-ranking poses (Supporting information Fig. S10). In one conformation PF-3845 is bended to interact with the additional pocket outside Phe pocket by π-π interactions. Another possibility is that PF-3845 could extend deep into the ATP pocket. Further work is needed to validate which binding mode is dominate.

### Co-crystal structure of PheRS-PF3845 and the role of a sulfate ion in the ligand binding

To illustrate the binding mode of PF-3845, we obtained the co-crystal structure of Mtb PheRS in complex with PF-3845 at a resolution of 2.3 Å. PF-3845 binds clearly in the amino acid pocket as well as the ATP pocket while the additional pocket is empty (Fig. 4C). An extensive hydrophilic and hydrophobic interaction network in two pockets can be found between PF-3845 and Mtb PheRS. Like the phenylsulfonamide group of GDI05-001, the 4-trifluoromethyl-2-pyridinyl group and phenyl ring of PF-3845 inserts deeply into the amino acid pocket. Two strong π-π stackings also can be seen between the phenyl rings of PF-3845 and F255/F257 residues of the PheRS (Fig. 4E). A hydrogen bond is formed between the amide of the trifluoromethylpyridinyl group and the main chain oxygen of G218. Two more hydrogen bonds are formed between the oxygen linker and the side chain of E217 and the main chain of F306. Unlike GDI05-001, the other end of PF-3845, including the 3-aminopyridine and piperidine groups, completely inserts into the ATP pocket (Fig. 4C and E). The carbonyl group of PF-3845 forms a hydrogen bond with the main chain amide of G309. The amide of 3-aminopyridine forms a water-mediated interaction with the side chain of E311 and the main chain of E311/R312. The 3-aminopyridine of PF-3845 also forms a strong π-π stacking with F213 and a π-cation interaction with R312. Surprisingly, a sulfate ion (SO_4_^2-^) is found near PF-3845 to bridge the interaction between the amide linker of PF-3845 and R201/R312 of the PheRS. Compared with the apo form, the side chain of R312 in the complex rotates nearly 90º to get closer to the compound. The SO_4_^2-^ most likely stabilizes R312, which in turn forms the π-cation interaction with the 3-aminopyridine. When this is compared with two other structures, hFARS2 complexed with Phe-AMP and *E. coli* PheRS (EcPheRS) complexed with Phe and AMP, the SO_4_^2-^ occupies an approximate space of AMP phosphate group in either case (Fig. 5). In the hFARS2-PheAMP co-crystal structure (32), the PheAMP forms hydrogen bond with R143 that is corresponding to R201 in Mtb PheRS. In the EcPheRS-AMP co-crystal structure, the AMP forms hydrogen bond with R301 corresponding to R312 in Mtb PheRS. Overall, the SO_4_^2-^ here engages R201 and R312 in the ATP pocket, and enables multiple interactions to facilitate PF-3845 binding into the ATP pocket of Mtb PheRS. Although SO_4_^2-^ has been found at the active site of several other aaRS structures, none of them contributes to the binding of a ligand as in the case of PF-3845 (33, 34).

**FIGURE 5.**
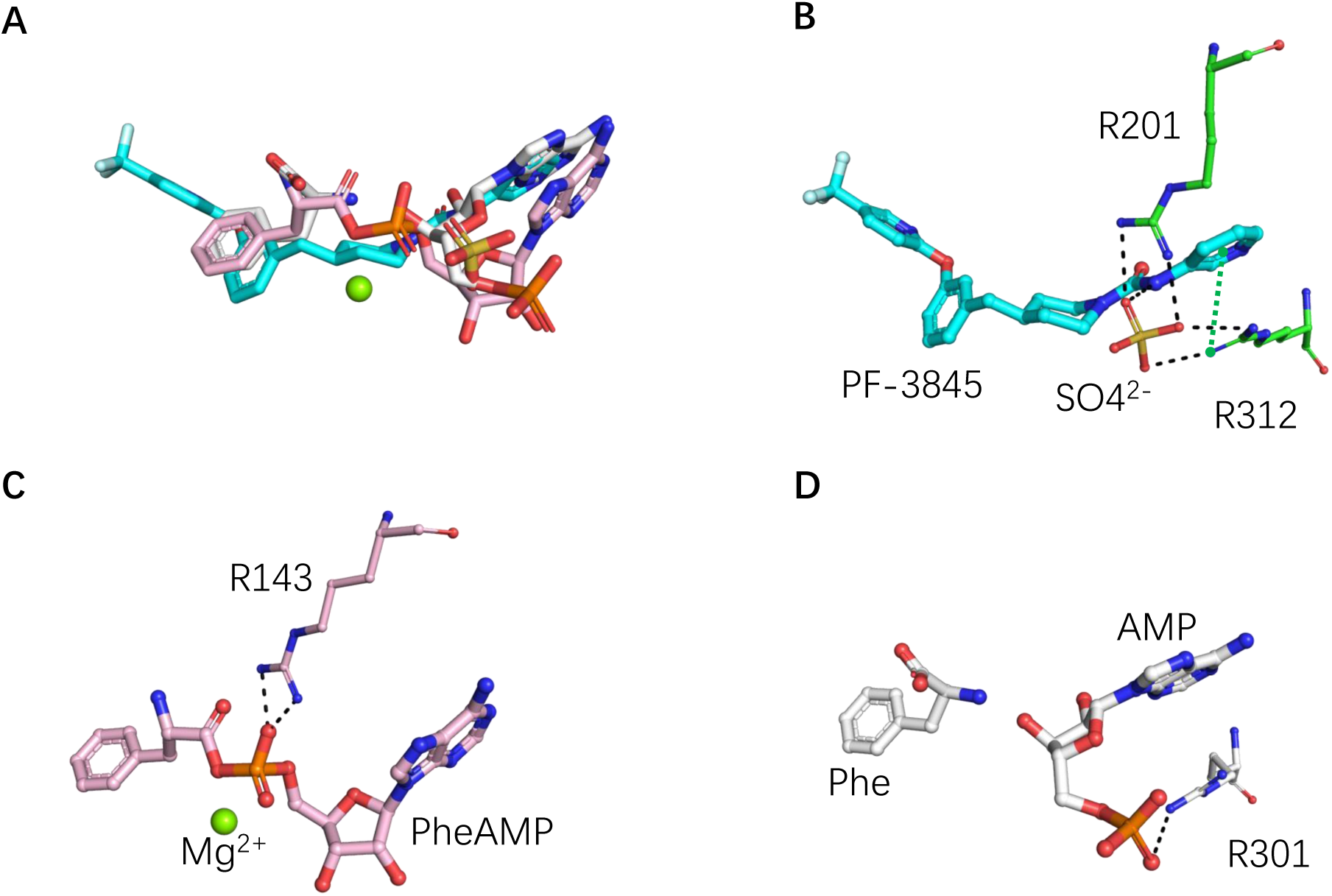
Sulfate ion stabilizes the interaction between PF-3845 and Mtb PheRS. A, structural comparison of the bound ligands of Mtb PheRS-PF-3845, hFARS2-PheAMP (PDB: 3CMQ) and EcPheRS-Phe-AMP (PDB: 3PCO). B, the SO_4_^2-^ interacts with the amide linker of PF-3845 and R201/R312 in MtbPheRS-PF-3845. C, the PheAMP interacts with R143 in hFARS2-PheAMP structure. D, the AMP interacts with R301 in EcPheRS-Phe-AMP structure.

To confirm the role of the SO_4_^2-^, we re-performed and analyzed the PF-3845 inhibition kinetics with 5 mM (NH_4_)_2_SO_4_ in the PheRS enzyme reaction. This time, PF-3845 clearly acted as an ATP competitive inhibitor (Fig. 3D), with the *K*_*i*_ calculated to be 0.73 ± 0.055 µM, which is lower than *K*_*i*_ 1.08 ± 0.245 µM with regard to ATP when SO_4_^2-^ was absent (Fig. 3B). The experiment demonstrated sulfate ion increased the binding of PF-3845, making it a true bi-substrate competitive inhibitor of Mtb PheRS.

### Cellular activity and target engagement of PF-3845

The MIC of PF-3845 against Mtb H37R_v_ measured with a microplate alamarBlue assay (MABA) (35), was ∼23.87 µM (n=3) compared to the MIC of GDI05-001 at 145 µM. It indicates PF-3845 has better permeability into Mtb cells than GDI05-001 even though its *K*_*i*_ to PheRS target is 9-fold higher. The cytotoxicity of PF-3845 or PF-04457845 (CC_50_), shown as 50% inhibition of the proliferation of mammalian cell lines, ranges from 16.9 to 25.8 µM (Supporting information Fig. S11), which is not surprising for a drug originally discovered as a human protein inhibitor. Next, we set to analyse if PF-3845 can engage PheRS target in Mtb H37R_v_. For this we used Mtb pheS-FDAS in which *pheS* was tagged to encode a C-terminal degradation tag. The IC_50_ of PF-3845 shifted down approximately 8-fold from 75 µM in WT H37Rv to 9.2 µM in the pheS-FDAS mutant, suggesting that PF-3845 can inhibit PheRS in Mtb (Fig. 6).

**FIGURE 6.**
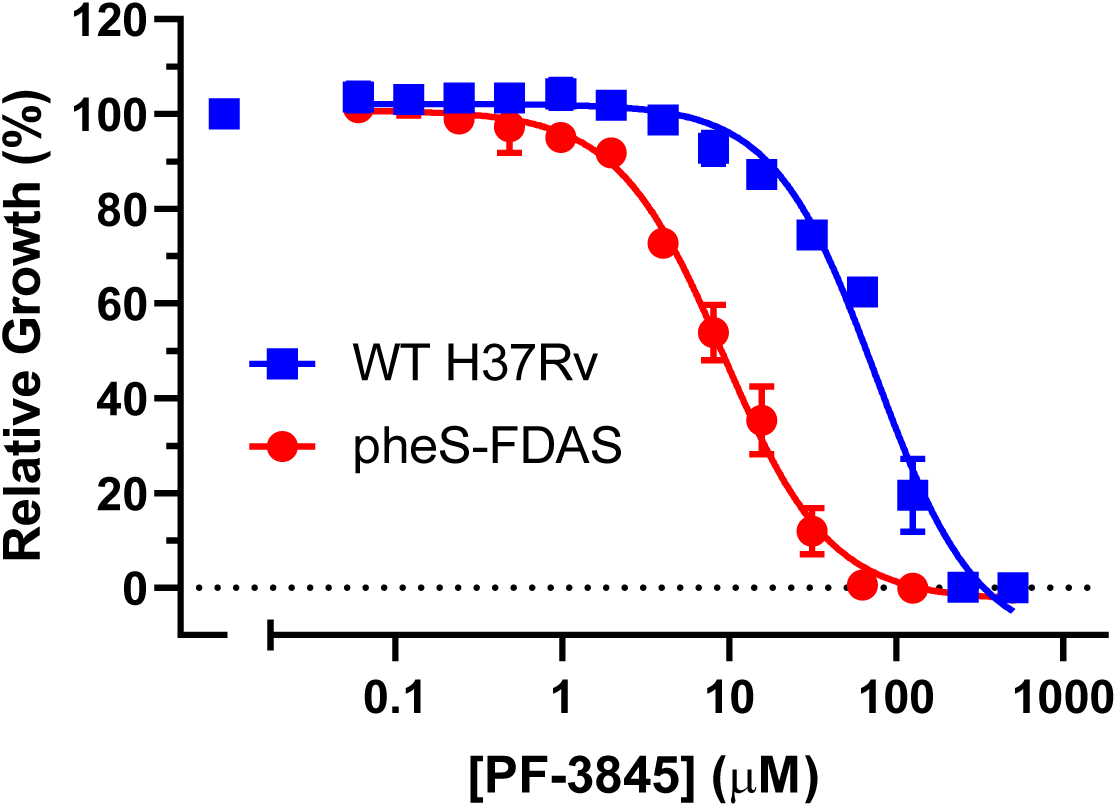
Susceptibility of WT H37Rv (blue squares) and pheS-FDAS (red circles) to PF-3845. Flag- and DAS-tagging PheS in strain pheS-FDAS derived from wide-type *Mycobacterium tuberculosis* H37Rv caused the IC_50_ of PF-3845 to shift down 8-fold compared to WT H37Rv. Results are representative of two independent replicates.

## Discussion

GDI05-001 belongs to the phenyl-thiazolylurea-sulfonamide class of PheRS inhibitors originally discovered in 2004 by researchers at Bayer Healthcare (12). Ten years later, an AstraZeneca pharmaceutical research group obtained the co-crystals of *Pseudomonas aeruginosa* PheRS in complex with GDI05-001 and identified an auxiliary pocket that is an extension of the phenylalanine binding pocket (16). It was argued that the auxiliary pocket is a liability for drug discovery because compound binding in the Phe pocket of bacterial PheRS leads to high screening hit rates, resistance frequencies, and elevated plasma protein binding. Further attempt to improve the physicochemical properties of inhibitors by exploring the hydrophilic ATP binding pocket resulted in very limited success. In this work, by a small-scale screening with a commercial library of 2184 bioactive compounds, we identified a completely different chemical scaffold, PF-3845, as a sub-micromolar inhibitor of Mtb PheRS. PF-3845 binds to Mtb PheRS in a way distinctive from GDI05-001. Most strikingly, PF-3845 extends into the ATP pocket of PheRS with the help of an inorganic sulfate ion. It represents the first non-nucleoside bisubstrate adenylation inhibitor of PheRS. Based on the 2.3 Å co-crystal structure, the trifluoromethylpyridinyl group on the west side of PF-3845 could be minimized to reduce its binding to the auxiliary pocket while the N-pyridine-carboxamide on the east is elaborated (Fig. 2B). This type of modification would overcome the shortfalls of the chemical scaffolds Bayer and AstraZeneca had previously pursued, turning up opportunities for design of TB drug candidate as well as new anti-Gram-negative lead. As the PF-04457845 has already shown a fantastic pharmacokinetic profile in rats and dogs as well as in vitro assays with human blood (36), other compounds derived from PF-3845 could also be promising clinical candidate for TB treatment.

Although the piperidine/piperazine urea group of PF-3845 occupies the ATP pocket, it doesn’t bind as deep as nonhydrolyzable phenylalanyl-adenylate analogue (PheOH-AMP) or Phe-AMP (37, 38). The latter forms multiple hydrogen bonds with a loop region. Two hydrogen bonds formed between PheRS and PF-3845 are through water or sulfate in buffer. The sulfate plays an import role in forming an interaction network including electrostatic, cationic-π, and hydrogen bonds that all together hold the piperidine carbamate inside the ATP pocket. Taking advantage of the available co-crystal complex structure, further optimization can be done rationally. For example, it could gain entropic contribution to the binding affinity if the compound is modified to replace the bridging water. Regarding the sulfate ion, one possible strategy is to grow a piece from PF-3845 to mimic its function for binding without losing cellular activity. Alternatively, no attempt to displace the sulfate ion could be made in design because it appears readily available due to a good number of sulfate transporters and sulfatases in mycobacteria (39).

PF-3845 does not inhibit either human cytoplasmic or mitochondrial PheRS in biochemical assays at concentrations up to 200 µM (Supporting information Fig. S6 and S12); the experimental data are supported by protein structural analysis. When comparing the catalytic site of Mtb PheRS with hFARS1 (40) and hFARS2 by structural superposition and sequence alignment, a major difference is found in the amino acid pocket (Supporting information Fig. S13). Residues F438 in hFARS1 and M258 in hFARS2, which form the pocket at the same position, are much larger than the corresponding residue V286 in Mtb PheRS, resulting in the smaller amino acid pocket of hFARs. Especially, F438 of hFARS1 reaches the position that could have steric crash with the thiazole ring of GDI05-001. GDI05-001 can therefore be effectively blocked from binding to human PheRSs. This analysis agrees with previous toxicity data (12, 16), and explains the selectivity and antibacterial potency of GDI05-001. Likewise, there is not enough space in hFARSs to accommodate PF-3845 since its two linked aromatic moieties take the same space as the thiazole ring of GDI05-001. Thus, the two hFARSs are not inhibited by PF-3845. Because PF-3845 was proven a covalent inhibitor of human FAAH, the cytotoxicity of PF-3845 most likely comes from inhibition of the hydrolase and/or other enzymes that have a nucleophilic group responsible for its molecular function. Medicinal chemistry effort is needed to change the carboxamide group in PF-3845 scaffold when it is modified toward a TB drug lead. Numerous PheRS enzymatic assays detailed in this work will assist the effort on track.

However, PF-3845 (PubChem CID: 25154867) has its own complexity. Its molecular weight of 456.5 g/mol and cLog*P* equivalent of 4.4, are at the upper limit of Pfizer’s rule of five that evaluates the drug-likeness of a compound based on orally active drugs for humans. A note of caution is the CC_50_ of PF-3845 to mammalian cells is even lower than its MIC against Mtb H37Rv, leaving it at a position unpropitious to become a TB drug candidate. Nevertheless, as tens of synthetic analogs of PF-3845 have been reported in literatures (29, 41, 42), the PF-3845 scaffold will serve as an excellent chemical probe to study the functions of bacterial and human PheRSs.

### Experimental Procedures

PheRS expression plasmids―The native sequences of Mtb *pheS* (NCBI Gene ID 885105) and *pheT* (NCBI Gene ID 885283) were synthesized by GenScript and co-cloned into the NdeI and HindIII sites of pET-30a expression vector. Between the two ORFs, a ribosome binding site (underlined) (…*pheS*…GGTGCCTAGTCTAGaaactaag<u>aaggag</u>atatacatATGGCCAGC…*pheT*…) was incorporated to increase the expression level of *pheT*. Only the C-terminal of PheT was designed to be fused to a His6 tag for co-purification of the two subunits. (Supporting information Fig. S1A).

Overexpression and purification of PheRS―The expression plasmid was transformed to BL21(DE3) competent cells and transformants were selected by antibiotic marker. Single colony was inoculated into 220 ml LB medium containing 20 μg/ml kanamycin or 100 μg/ml ampicillin and the culture was incubated for 16 hours at 37 °C. 9-ml fresh culture was used to inoculate 1 L LB broth medium (total 24 L for one preparation) and grown at 37 °C until the OD_600_ reached 0.7-0.9. The culture was cooled down to 16°C, then IPTG was added to the final concentration of 0.3 mM. After 18 hours incubation at 16 °C, cells were harvested by using centrifuge (3800 g, 15 min, 4°C) and resuspended with cell harvesting buffer (25 mM Tris pH8.0, 150 mM NaCl, 10mM imidazole pH8.0, 1 mM DTT and EDTA-free protease inhibitor tablet (Roche)). Resuspended cells were divided into 2 portions and handled independently to lyse through cell disrupter (700-800 bar, 5 cycles) followed by 3-minute sonication (60% amplitude). Cell debris were removed through centrifugation (24000 g, 80 minutes) followed by filtration against 0.22 μM PVDF membrane (Millipore), then the supernatant was loaded onto HisTrap-FF (GE Healthcare, 1 column volume (CV) = 5 ml, 2 columns in tandem). The flow rate was 2.5 ml/min. After washing 20 CV with washing buffer (25mM Tris pH 8.0, 150mM NaCl, 20mM imidazole pH8.0, 1 mM DTT), fractions were collected with gradient elution buffer (25 mM Tris pH 8.0, 150 mM NaCl, 20-300 mM Imidazole pH8.0, 1 mM DTT). Based on samples examined with SDS-PAGE and stained with Coomassie blue, the proper fractions were combined and applied to anion exchange purification with HiTrap Q-HP (GE, 1CV = 5 ml, 2 columns in tandem). After washing 12 CV with washing buffer (25 mM Tris pH 8.0, 10 mM NaCl, 1 mM DTT), fractions were collected with gradient eluted buffer (25 mM Tris pH8.0, 20-500 mM NaCl, 1 mM DTT). Proper fractions based SDS-PAGE analysis were collected and concentrated to about 2ml and injected to Superdex-200 increase (flow rate: 0.3 ml/min, ∼1.4 Mpa, GE) for further purification. The proper fractions were pooled together (25 mM HEPES pH7.5, 150 mM NaCl, 5 mM MgCl_2_ and 5% glycerol) and concentrated to about 2 ml using Amicon-centrifugal filter (Millipore). UV_280_ absorbance of PheRS was measured on NanoDrop OneC (Thermo). Extinction coefficient of PheRS (ε = 111840 M^-1^ cm^-1^) was calculated by entering alpha-beta heterodimer sequences into protparam program on ExPASy webserver. PheRS used in non-radioactive assays was pooled with pure fractions of multiple rounds of preparation and the protein concentration was calculated by Beer’s Law. It was dispensed into 50 μl aliquots and stored at −80°C for biochemical assay and crystallization trials.

tRNA expression plasmid―Sequence of Mtb-tRNA_TGG_^Phe^gene with T7 promoter (underline) at the 5’-terminal and CCA at the 3’-terminal (<u>taatacgactcactata</u>GGCCAGGTAGCTCAGTCGGTATG-AGCGTCCGCCTGAAAAGCGGAAGGTCGCGGTTCGATCCCGCCCCTGGCCAcca) was synthesized (GenScript). An EcoRI site was designed upstream of T7 promoter and a BamHI site was downstream of the tRNA gene. The fragment was cloned into the EcoRI and BamHI sites of pTrc99a vector (Addgene).

Overexpression and partial purification of tRNA^Phe^―The expression plasmid was transformed to BL21(DE3) competent cells. After incubation overnight on Ampicillin containing LB agar, single colony was inoculated to 100ml LB broth medium containing 100 μg/ml Ampicillin and incubated for 16 hours at 37 °C. 10 ml fresh culture was used to inoculate 1L LB broth medium (total 10L for one preparation) and grown at 37 °C until the OD_600_ reached 0.6-0.8, then IPTG was added to final concentration of 0.4 mM. After 16 hours incubation at 37°C, cells were harvested by using centrifuge (3800 g, 15 min, 4°C) and resuspended in cell lysis buffer (100 mM Tris pH7.0, 20 mM MgCl_2_; 30 ml per 1 L culture). Then equal volume of water-saturated phenol pH4 was added and fully mixed at room temperature for one hour. After centrifugation (14000 g, 30 min, 25 °C), the upper aqueous layer was carefully transferred to be mixed with equal volume of chloroform. The mixing was gently performed, and the mixture was let stand for a while. After short centrifugation, carefully transfer the upper layer into multiple new tubes. Triple volumes of pre-cold ethanol were added in and nucleic acid was precipitated at −20 °C for more than two hours. Supernatant was discarded after centrifugation (14000 g, 30 min, 4 °C). A pellet was dissolved in 10 ml deacylation buffer (500 mM Tris pH9.0) and incubated at 37 °C for one hour. Nucleic acid was afterward precipitated as before, and a pellet was washed two times with 75% pre-cold ethanol by centrifugation, dried in air and re-dissolved in 10 ml DEPC-treated water. The dissolved nucleic acid was mixed with 1 M MOPS pH7.0 buffer to a final concentration of 0.1 mM. One Q-2500 column (Qiagen) (for every 2 L starting culture) was pre-equilibrated with 50 ml equilibration buffer twice (50 mM MOPS pH7.0, 15% isopropanol, 1% Triton-X100). The nucleic acid was bound onto the column and washed four times with 50 ml washing buffer (50 mM MOPS pH7.0, 200 mM NaCl) by gravity flow. Then, each column was washed with 10 ml elution buffer once (50 mM MOPS pH7.0, 650 mM NaCl), followed by at least 2×10 ml the same buffer that eluted mainly tRNAs. The elution fractions could be examined by TBE Urea gel (Novex) and pure tRNA factions were precipitated by adding 1/10 volume of 3 M sodium acetate and equal volume of isopropanol for two hours at −20°C. The pellet was washed twice with 75% pre-cold ethanol by centrifugation, air dried and dissolved in tRNA storage buffer (50 mM HEPES pH7.4, 20 mM MgCl2, 100 mM KCl) to final concentration of 10mg/ml. It was dispensed to 100 μl aliquots and stored at −20°C.

Pyrophosphate (PPi) production assay―The assay was based on modification of EnzChek™ Phosphate assay kit (Thermo). Several aaRS works reported using this assay (3, 27, 28, 43). Our assay was performed in Corning 384-well flat clear bottom black microplate (cat #3764) and the reaction plates were read on EnVision (PerkinElmer). The final concentration of each component in the reaction buffer is 50 mM HEPES pH7.4, 100 mM NaCl, 10mM MgCl2, 1mM TCEP, 0.1 mg/ml BSA, 0.01% BriJ-35 (Sigma). In IC_50_ determination, the reaction was started with 20 μl volume in a well containing 50 nM Mtb PheRS, 0.5 mM ATP, 50 μM L-phenylalanine, 0.1 mg/ml tRNA^Phe^, 0.05 mM MESG, 0.5 u/ml PPase (Sigma), 0.1 u/ml PNPase (Thermo). Absorbance at 360nm was read every minute for 30 minutes. The PPi standard curve was made with 0.05 mM MESG, 0.5 u/ml PPase, 0.1 u/ml PNPase and 1.25 to 20 μM sodium pyrophosphate, the reactions was incubated for 15 minutes before read at 360 nm. Readout was converted to PPi production using a standard curve. GraphPad Prism 10.0 was used for data analysis.

ATP consumption assay and HTS screening―This assay used for HTS screening was based on Kinase-Glo luminescent Kit (Promega). The application with aaRS was previously descried (28, 44). Screening was carried out in Corning 384-well white flat bottom microplates (cat #3570). The final concentrations in the reaction are 25 mM HEPES pH7.4, 140 mM NaCl, 40 mM MgCl2, 30 mM KCl, 1 mM TCEP, 0.1 mg/ml BSA, 0.004% Tween-20. Compounds from Selleck-2148 bioactive library (Selleckchem) 10 mM stock were prepared in assay plate using Echo 550 (Labcyte) to final concentration of 50 µM. The 50-nL compound volume is negligible in each reaction. DMSO was added into negative control wells and tool compound GDI05-001 in positive control wells. 5μl Mtb PheRS in HEPES buffer was added to 100 nM using Multidrop dispensers (Thermo) and the compound-enzyme mixture was incubated for 30 minutes. Then 5-μl substrate mixture (1 μM ATP, 20 μM L-phenylalanine and 0.1 mg/ml tRNA^Phe^) was added to start the reaction at 37°C and incubated for 2 hours. After cooling down the plates, 10-μl diluted Kinase-Glo Max reagent (1/50 diluted with buffer 50 mM Tris pH7.5, 5% glycerol) was added and incubated for 15 minutes. Luminescence was recorded using SpectraMax M5 (Molecular Devices).

IC50 determination―Each selected compound was prepared using Echo 550 in 384-well plates in a series of 10 concentrations. DMSO wells were negative control and no enzyme wells were positive control. For assay with Mtb PheRS, the same method was used as above. For hFARS1, 50 nM enzyme, 2 μM ATP, 20 μM L-phenylalanine and 1 mg/ml yeast total tRNA (Roche) was incubated with compound at 37 °C for one hour. For hFARS2, the same method as hFARS1 except that *E. coli* total tRNA (Roche) was used at 0.5 mg/ml. Plates were processed for luminescence as above. In data analysis with the Graphpad Prism. The enzyme remaining activity was calculated as:

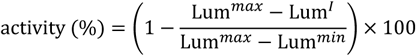

Lum^*max*^ was luminescence value without PheRS in reaction, Lum^*min*^ was the value from DMSO control well and Lum^*I*^ was from inhibitor well. The activity (%) was plotted against inhibitor concentration. To calculate IC50, the resulted dose-response curve was fit to Equation 1, where H was the Hillslope factor.

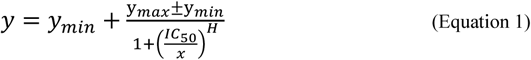

Mode of inhibition (MOI)― By the PPi production assay, 50 nM of MtbPheRS with two saturating substrates and one variable substrate, were used in the measurement of steady-state enzyme kinetics at a series of fixed concentrations of PF-3845 (MCE MedChemExpress). Lineweaver-Burk plots were fitted for modelling of inhibition (competitive, non-competitive, mixed or uncompetitive). To calculate the *K*_i_ for competitive inhibition, the plots with PPi production rate against substrate concentration were fit to:

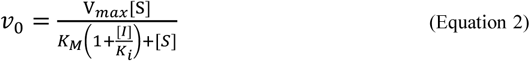

To calculate the *K*_i_ for mixed inhibition, the plots were fit to:

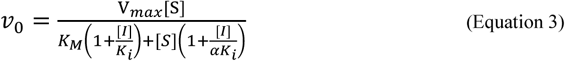

Crystallization, data collection and structure determination― Mtb PheRS was incubated with GDI05-001 or PF3845 at a molar ratio of 1:5 for 4 hours. 5 mg/ml of the apo form and two complexes were crystallized separately by sitting drop vapor diffusion method at 18 °C. The best crystals were obtained by seeding with well buffer containing 0.1 M HEPES pH 7.7, 1.5 M (NH_4_)_2_SO_4_, 0.2% polyethylene glycol (PEG) 3350. The cryo-protectant solution contained 0.1 M HEPES pH 7.7, 1.5 M (NH_4_)_2_SO_4_, 0.2% polyethylene glycol (PEG) 3350, 25% Glycerol. X-ray data were collected at beamline BL19U in Shanghai Synchrotron Radiation Facility (SSRF) at 100K. Data integration and scaling were performed using HKL2000. These crystals belong to the space group C2221, contain one PheRS heterodimer per asymmetric unit. The apo form structure was determined by molecular replacement with the Phaser-MR module in Phenix using *E. coli* PheRS (PDB code: 3PCO) as a search model. The model adjustment and refinement were performed using COOT and Phenix. Structures of the two liganded TbPheRS complexes were determined using the coordinate of the apo form structure. The final refinement statistics are summarized in Table 1. All figures were prepared using Pymol. The relevant coordinates and structure factors have been deposited in PDB with accession codes 7DAW, 7DB7 and 7DB8.

**TABLE 1.** Data-collection, phasing and refinement statistics.

| **Data collection** | MtbPheRS (apo form) | MtbPheRS-GDI05-001 | MtbPheRS-PF-3845 |
| --- | --- | --- | --- |
| Space group | C2221 | C2221 | C2221 |
| Resolution (Å)<sup>a</sup> | 50.0-2.83 (2.83-2.87) | 50.0-2.71 (2.71-2.75) | 50.0-2.30 (2.30-2.34) |
| Cell dimensions |  |  |  |
| a, b, c (Å) | 267.00, 298.91, 65.94 | 268.38, 298.80, 65.71 | 278.01, 297.81, 65.90 |
| $\alpha$, $\beta$, $\gamma$ (°) | 90, 90, 90 | 90, 90, 90 | 90, 90, 90 |
| Unique reflections | 59,426 (2,275) | 72,740 (3,594) | 121,928 (6,050) |
| I/σI | 10.86 (1.67) | 10.42 (3.26) | 19.0 (4.1) |
| Completeness (%) | 93.5 (71.5) | 99.9 (100.0) | 99.9 (100.0) |
| R<sub>merge</sub> (%) | 17.4 (45.5) | 13.3 (76.6) | 13.8 (68.1) |
| CC1/2 (%) | 96.4 (84.0) | 98.5 (85.1) | 99.4 (95.3) |
| Wilson B factor (Å<sup>2</sup>) | 59.36 | 45.37 | 29.83 |
| **Refinement** |  |  |  |
| R<sub>work</sub> (%) | 20.00 | 19.20 | 16.83 |
| R<sub>free</sub> (%) | 24.02 | 21.76 | 18.65 |
| Average B factors (Å<sup>2</sup>) | 61.02 | 43.58 | 35.26 |
| Root mean square deviations |  |  |  |
| Bond lengths (Å) | 0.009 | 0.009 | 0.008 |
| Bond angles (°) | 1.109 | 1.11 | 0.997 |
| Ramachandran plot |  |  |  |
| favored (%) | 94.26 | 95.27 | 96.99 |
| allowed (%) | 5.47 | 4.37 | 2.91 |
| outliers (%) | 0.27 | 0.36 | 0.09 |

a. Numbers in parentheses are for the highest resolution shell.

## Acknowledgements

This work is supported by Bill & Melinda Gates Foundation and GHDDI management. The authors thank the staff of beamline BL19U1 at Shanghai Synchrotron Radiation Facility for access and help with the data collection. Dr. Wen Xiong participated in planning the project. S.C. and H.W. appreciate the help provided by Prof. Babak Javid’s lab in Tsinghua University and Prof. En-Duo Wang’s lab in CAS Shanghai Institute of Biochemistry and Cell Biology.

## ^1^Abbreviations and nomenclature

PheRS: bacterial phenylalanyl-tRNA synthetase
FARS: human phenylalanyl-tRNA synthetase
Phe: phenylalanine
ATP: adenosine triphosphate
AMP: adenosine triphosphate
PPi: pyrophosphate
PPase: pyrophosphatase
Pi: phosphate
MESG: 2-amino-6-mercapto-7-methylpurine ribonucleoside
AMMP: 2-amino-6-mercapto-7-methylpurine.

## Supporting Information

**Figure S1.**
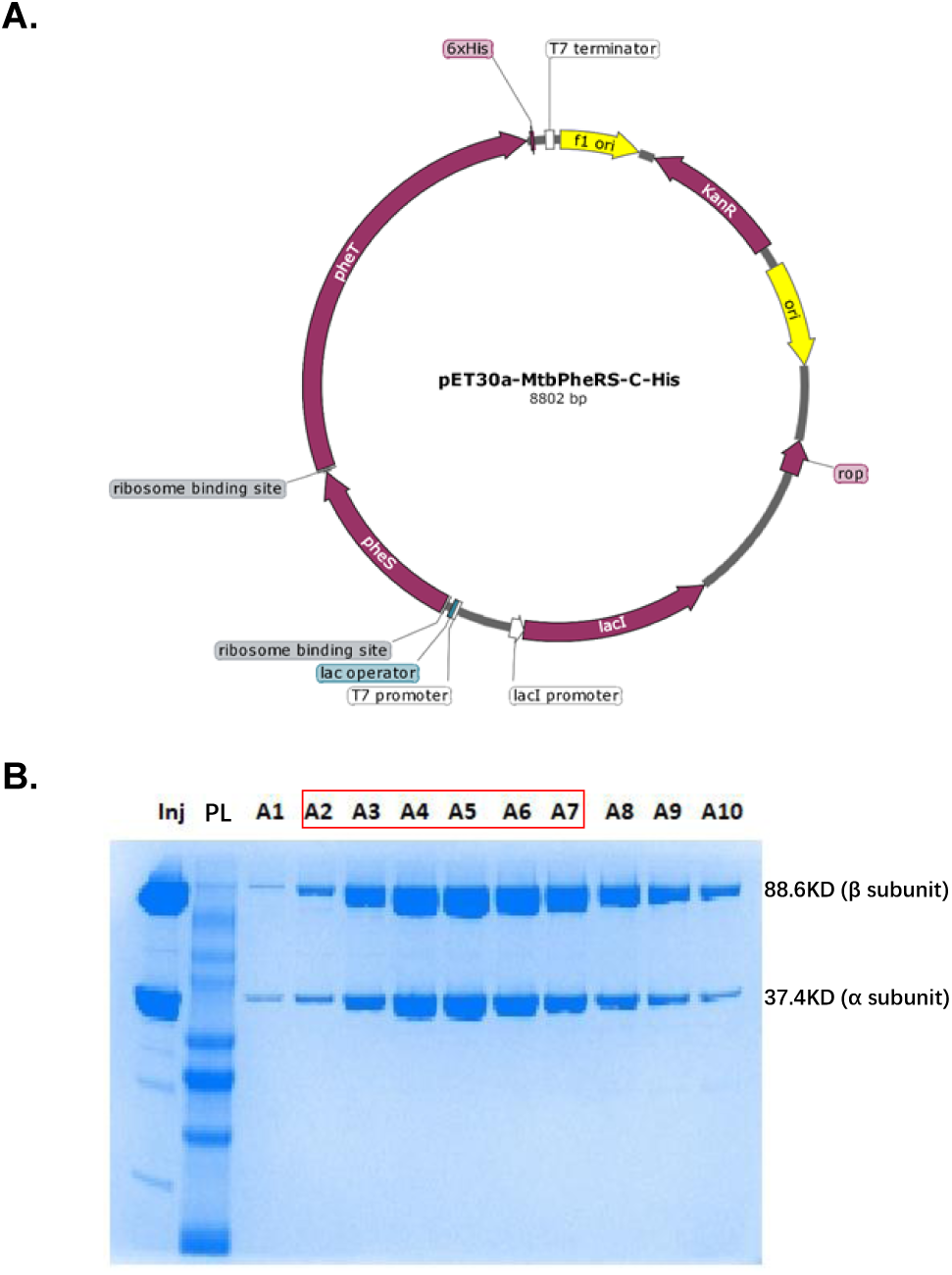
Plasmid map for PheRS overexpression in *E. coli* (A) and the purified PheRS shown by SDS-PAGE electrophoresis (B). The fractions (A1-A10) were from the last-step size exclusion column. Inj, sample before injecting into the column; PL, protein ladder. Red box indicates the collected fractions of PheRS.

**Figure S2.**
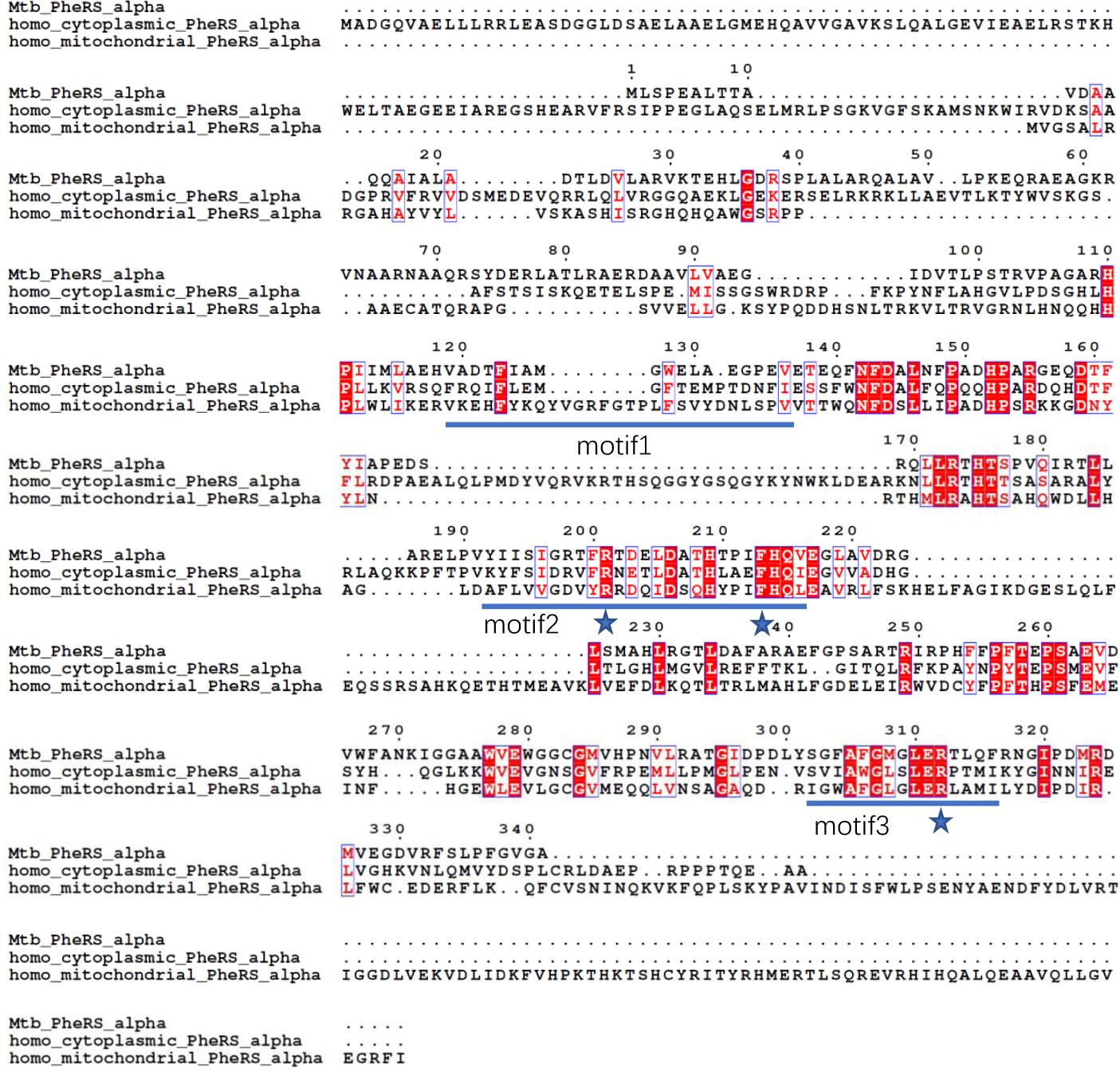
Alignment of PheRS α subunits and identification of motifs. Motif 1 forms part of the dimer interface and helps orient the motif 2 and 3 active site loops. Motif 2 participates in the binding of ATP, amino acid substrate, and the acceptor end of tRNA, while motif 3 interacts with ATP substrate. The highly conserved arginine and phenylalanine of motif 2, and the arginine of motif 3 marked by asterisks have catalytic roles.

**Figure S3.**
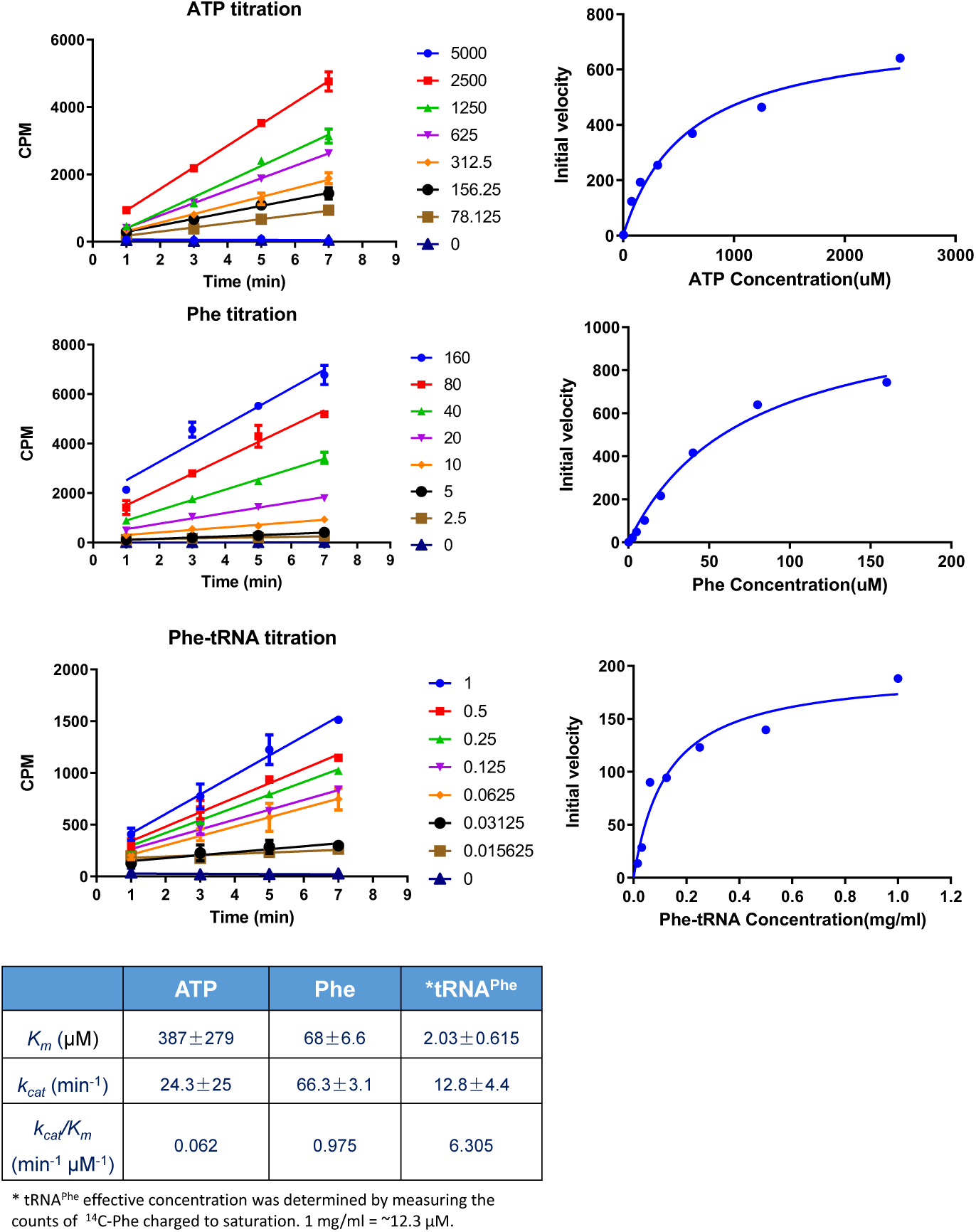
Michaelis-Menten kinetic parameters of Mtb PheRS determined by the aminoacylation assay. It is based on retention of L-[^14^C(U)]-phenylalanine charged tRNA on GF/B fiberglass filter.

**Figure S4.**
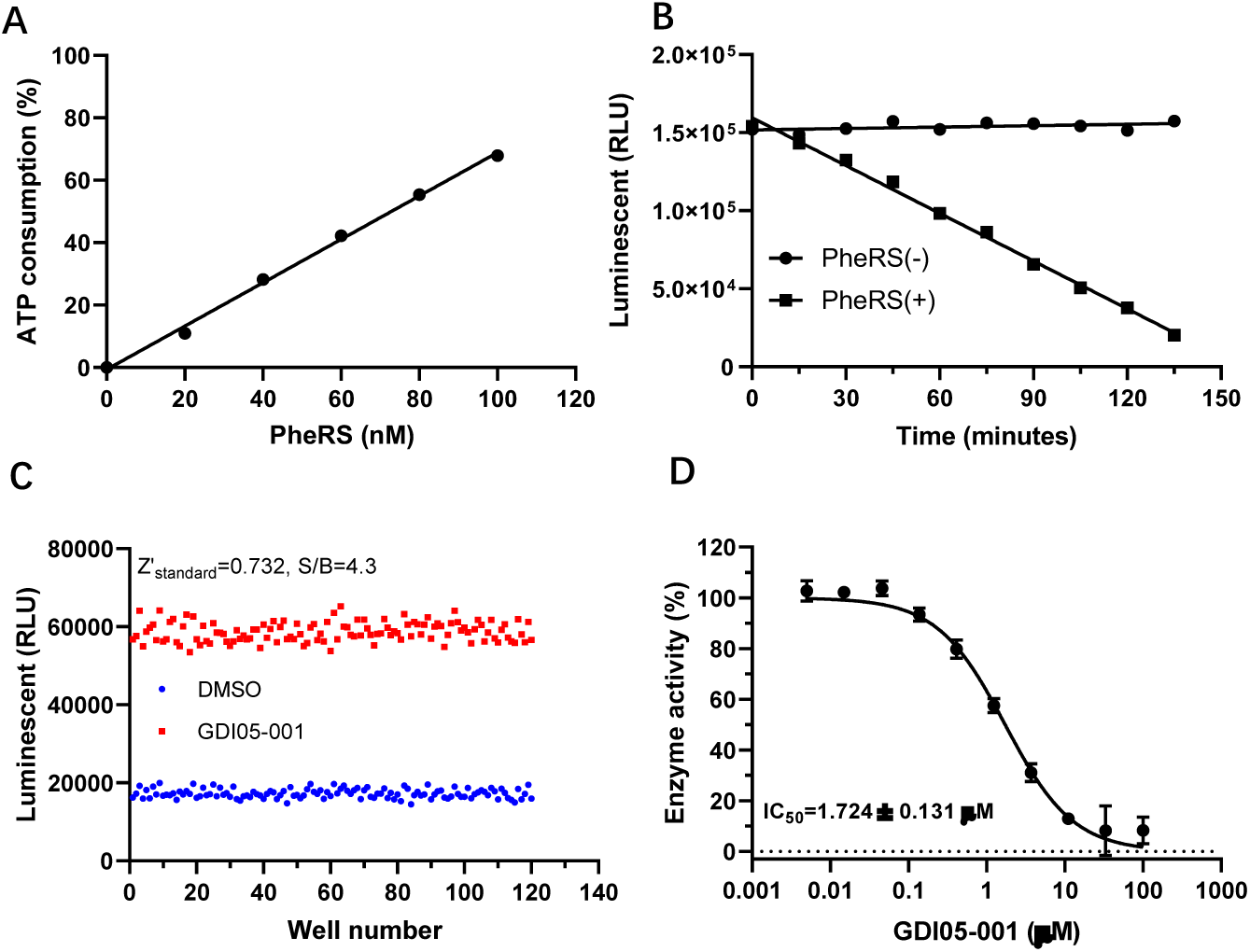
Development of HTS using ATP consumption assay (Kinase-Glo) and HTS optimization. A, Linearity of ATP consumption with respect to Mtb PheRS concentration. B, Linearity with respect to time with or without Mtb PheRS. C, Z’ factor of HTS by using ATP consumption assay. D, IC_50_ of tool compound GDI05-001 against Mtb PheRS.

**Figure S5.**
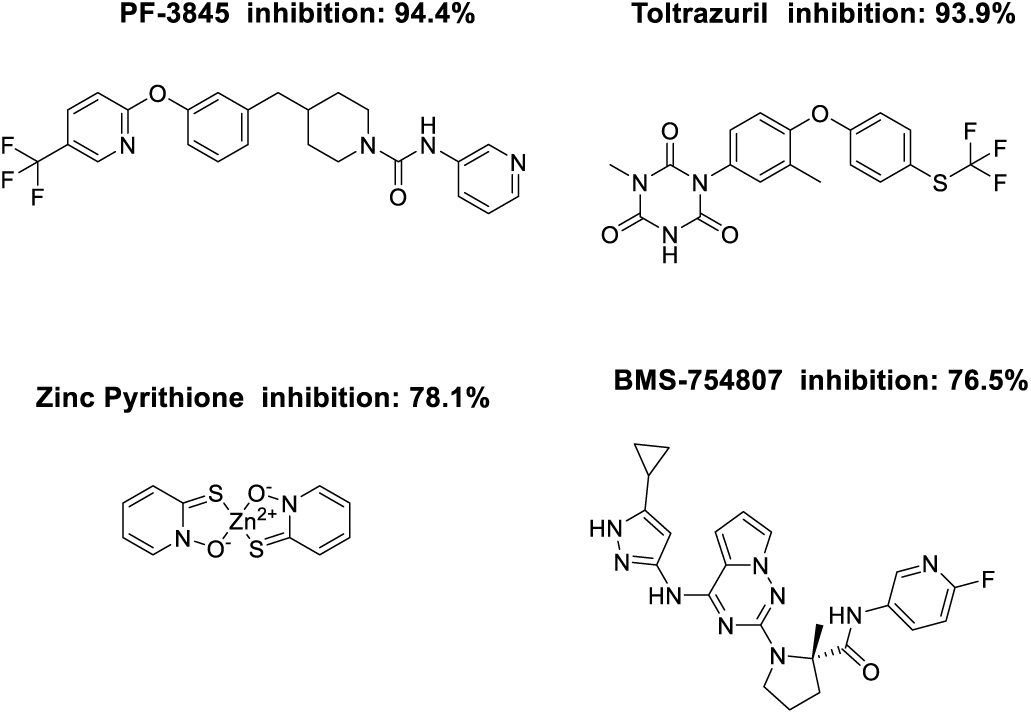
Primary hits of the Mtb PheRS HTS.

**Figure S6.**
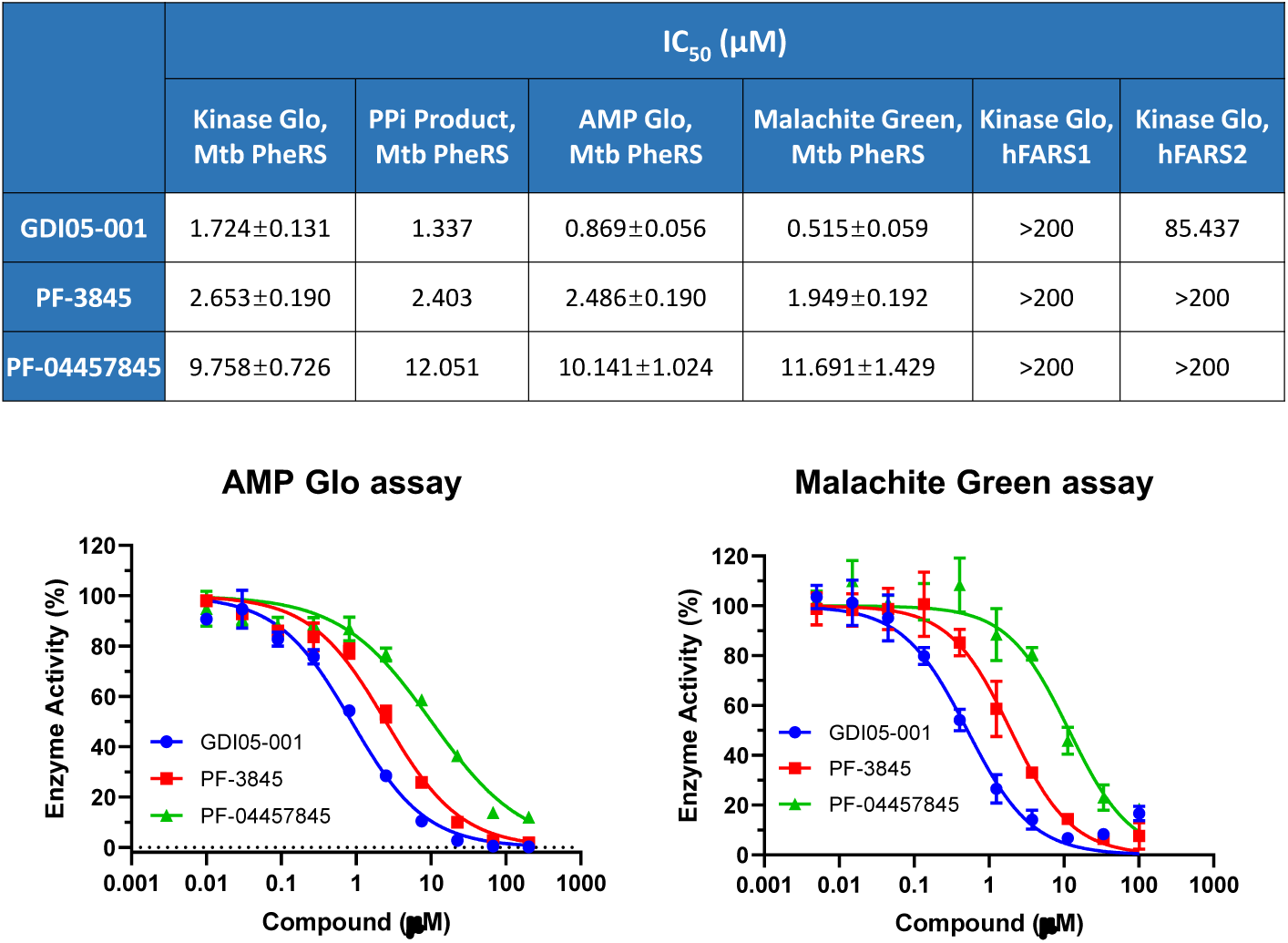
IC_50_s of compounds against Mtb PheRS or hFARSx determined by different other assays. Kinase-Glo and PPi production assays are mentioned in the text. AMP-Glo assay measures AMP production with an AMP-Glo kit (Promega). Malachite Green assay is an end-point assay measuring Pi production, previously applied to aaRS (ref. 30).

**Figure S7.**
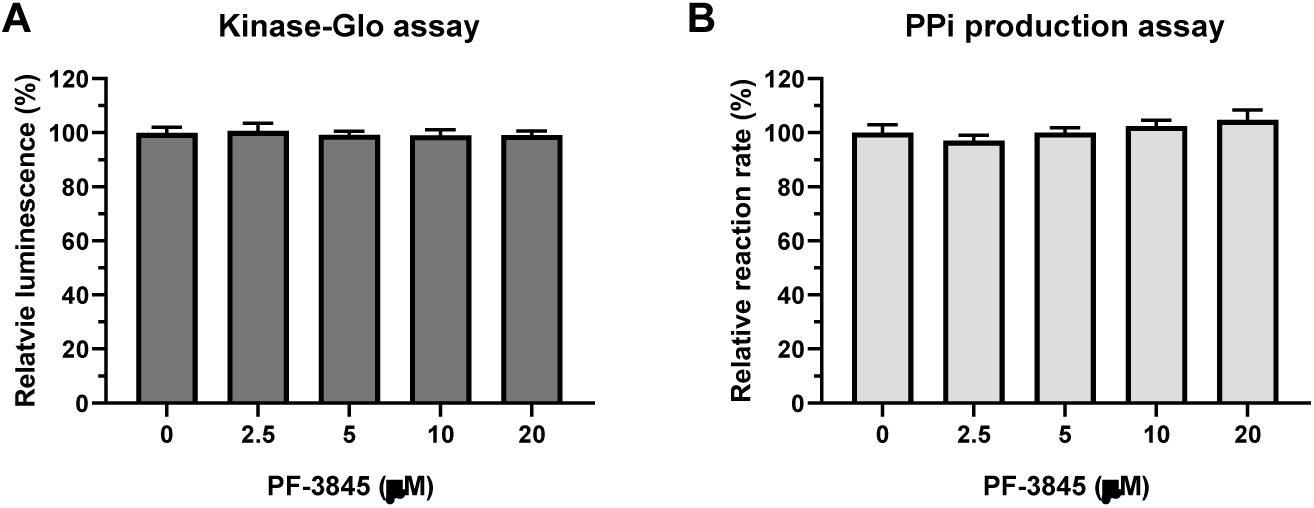
PF-3845 did not inhibit the coupled reactions in Kinase-Glo (A) and PPi production (B) assays. The two assay reactions were set up and monitored as those in IC_50_ determination except that Mtb PheRS was omitted.

**Figure S8.**
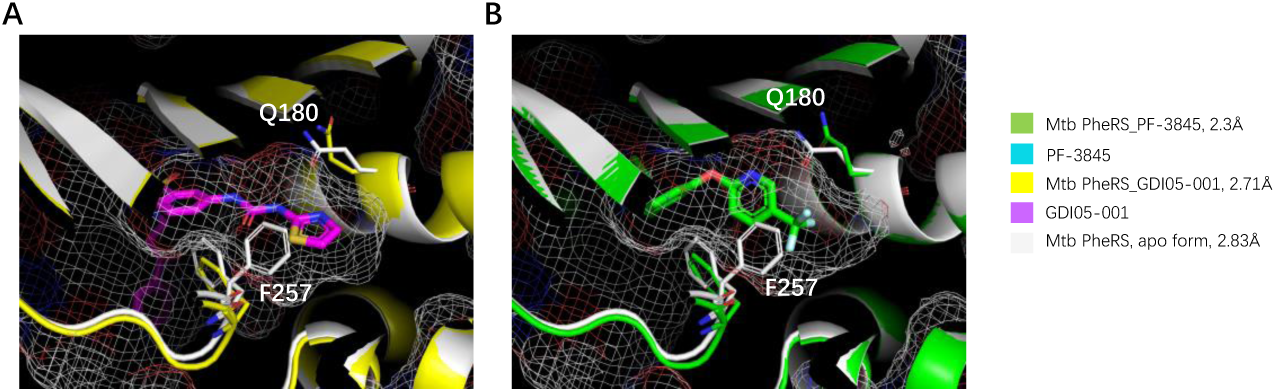
F257 and Q180 rotation creates a deeper amino acid pocket of Mtb PheRS. The side chain rotation of F257 and Q180 contributes to a much deeper amino acid pocket in the complexed structure than that in apo form structure. The amino acid pockets are indicated by mesh. A. Superposition of apo and GDI05-001 complexed structures. B. Apo and PF-3845 complexed structures.

**Figure S9.**
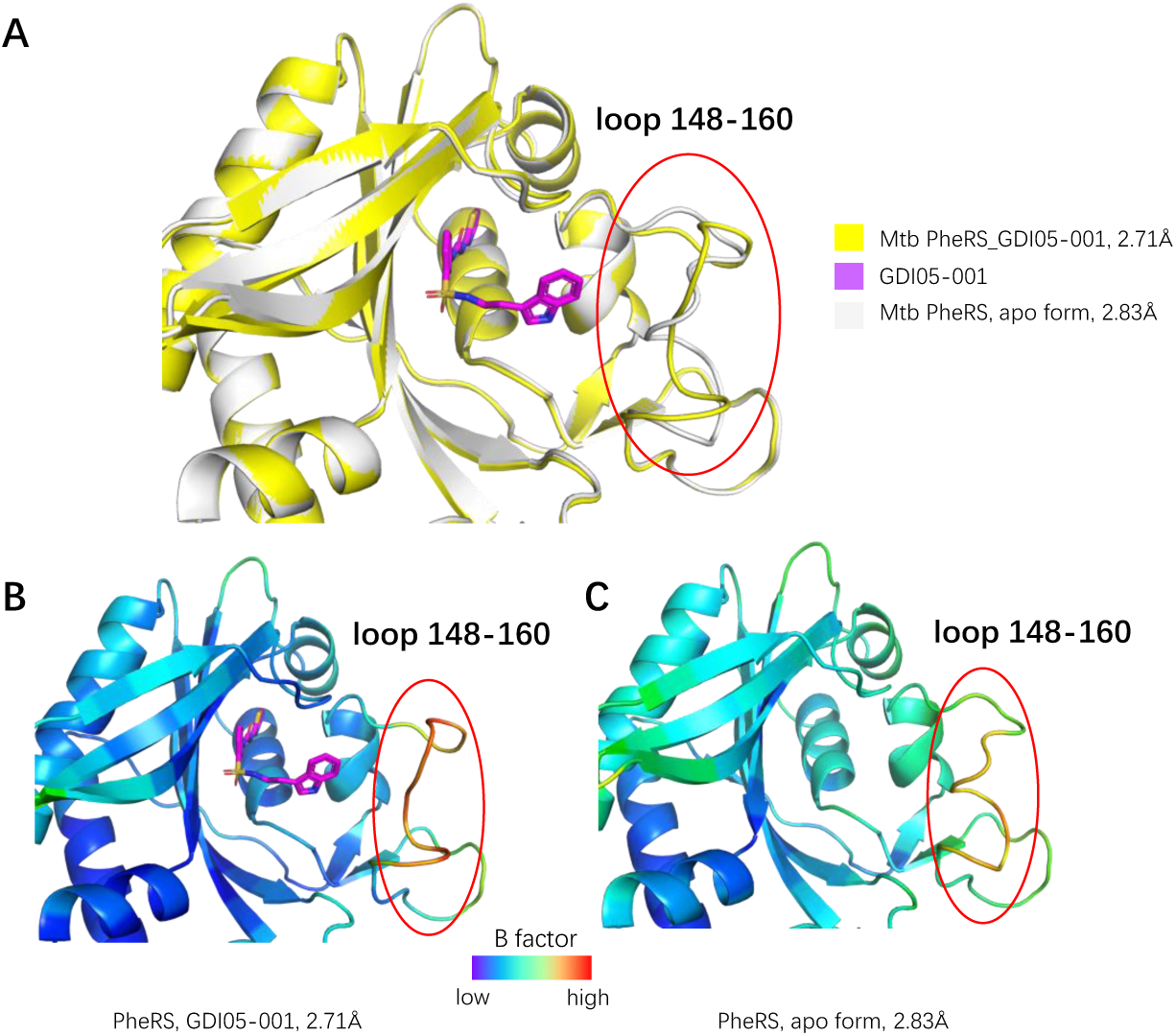
GDI05-001 binding leads to conformational change of the loop 148-160 of α subunit of Mtb PheRS. A, structure comparison of the α subunits of Mtb PheRS apo form and in complex with GDI05-001. B-C, structure colored by B factors. Compared to apo structure, the loop 148-160 of the GDI05-001 bound structure undergoes a conformational change, which has much higher B factors than the apo structure.

**Figure S10.**
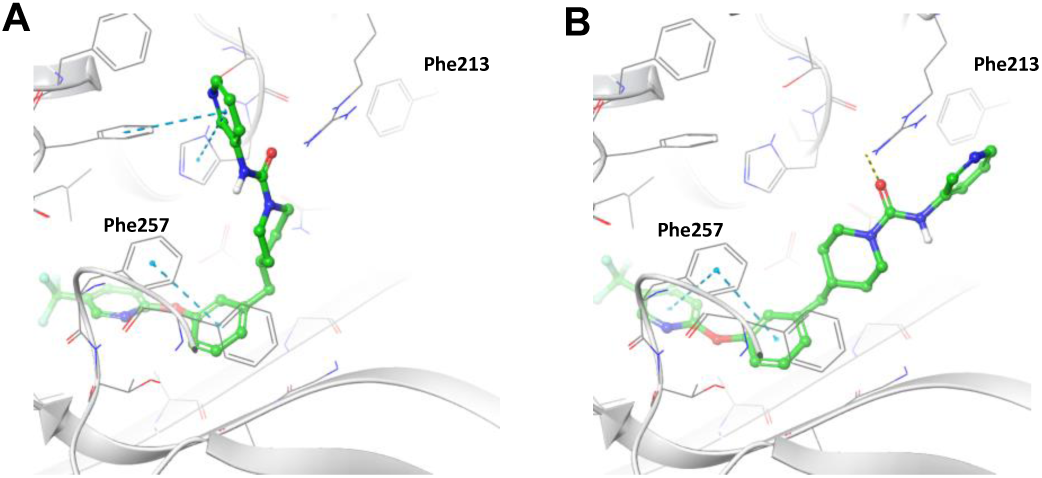
Two putative binding conformations of PF-3845 generated by Orbital docking platform (Accutar Biotech). A, PF-3845 is modelled into the substrate Phe and additional pockets. B, PF-3845 is modelled into both Phe and ATP binding pockets.

**Figure S11.**
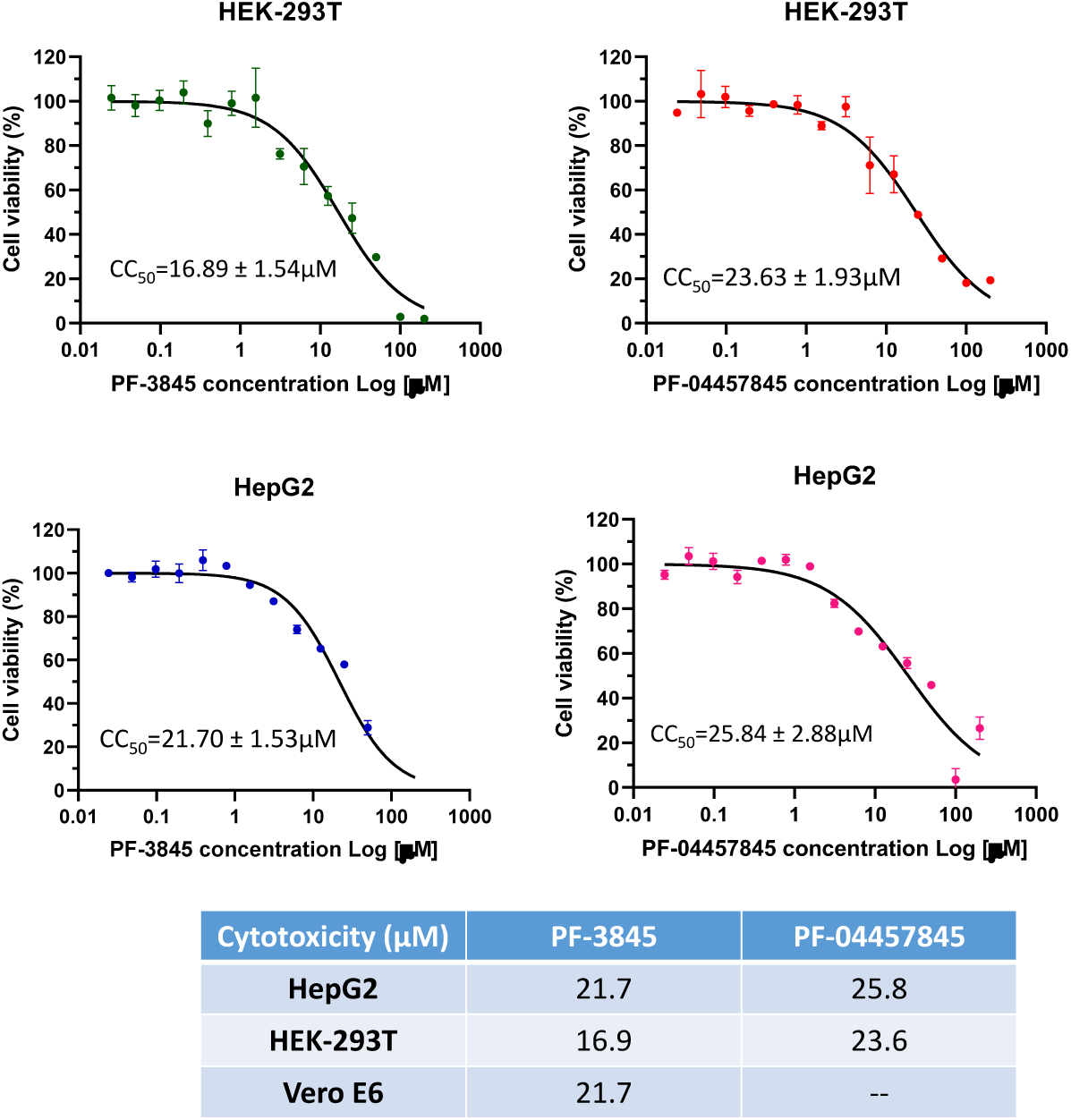
Cytotoxicity of PF-3845 and PF-004457845 against mammalian cell lines. HepG2 is an immortal cell line derived from human liver cells. HEK-293T is a human embryonic kidney 293 cell line. Vero E6 is a cell line derived from African green monkey kidney epithelial cells.

**Figure S12.**
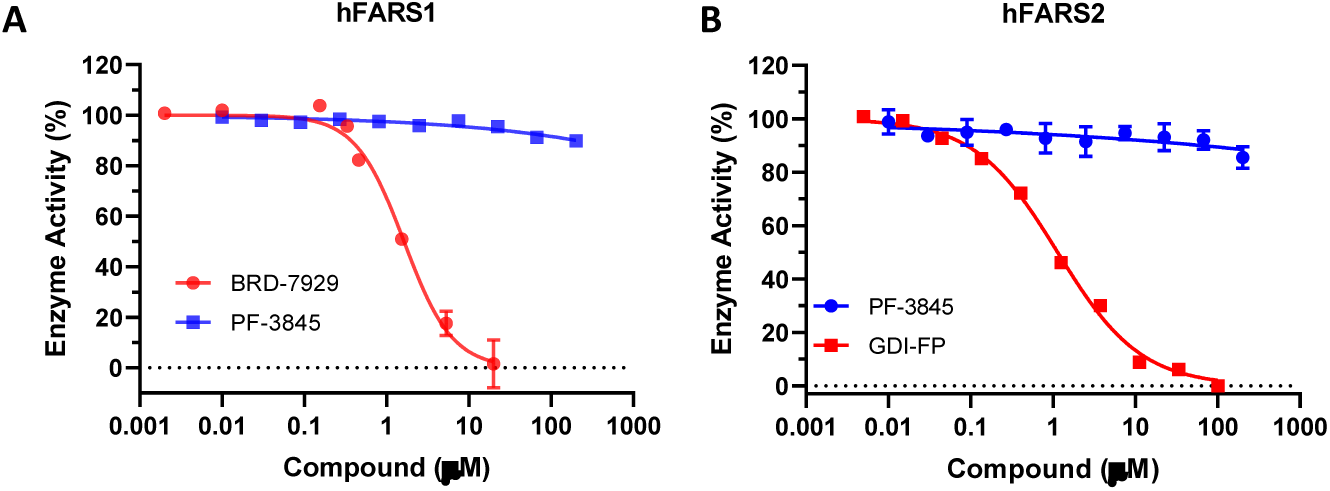
Analysis of potential effect of PF-3845 against human PheRSs, cytoplasmic hFARS1 (A) and mitochondrial hFARS2 (B). Compound BRD-7929, known to have a level of inhibitory effect on eukaryotic PheRS (ref.21), was used as positive control in the hFARS1 experiment. Compound GDI-FP undisclosed was used as positive control in the hFARS2 experiment. The same strategy for cloning Mtb PheRS was used to synthesize and construct expression plasmid for human cytoplasmic PheRS (hFARS1, Gene IDs 2193 and 10056). Gene for human mitochondrial PheRS (hFARS2, Gene ID 10667) did not include mitochondria signal peptide sequences (37 amino acids at the N-terminal), and was cloned into the NdeI and SalI sites of pET-21a vector, which gives a fusion protein with C-terminal His6-tag. Genes for the two human PheRSs were codon optimized for heterologous expression in *E. coli*. The protein purification steps were very similar to Mtb PheRS.

**Figure S13.**
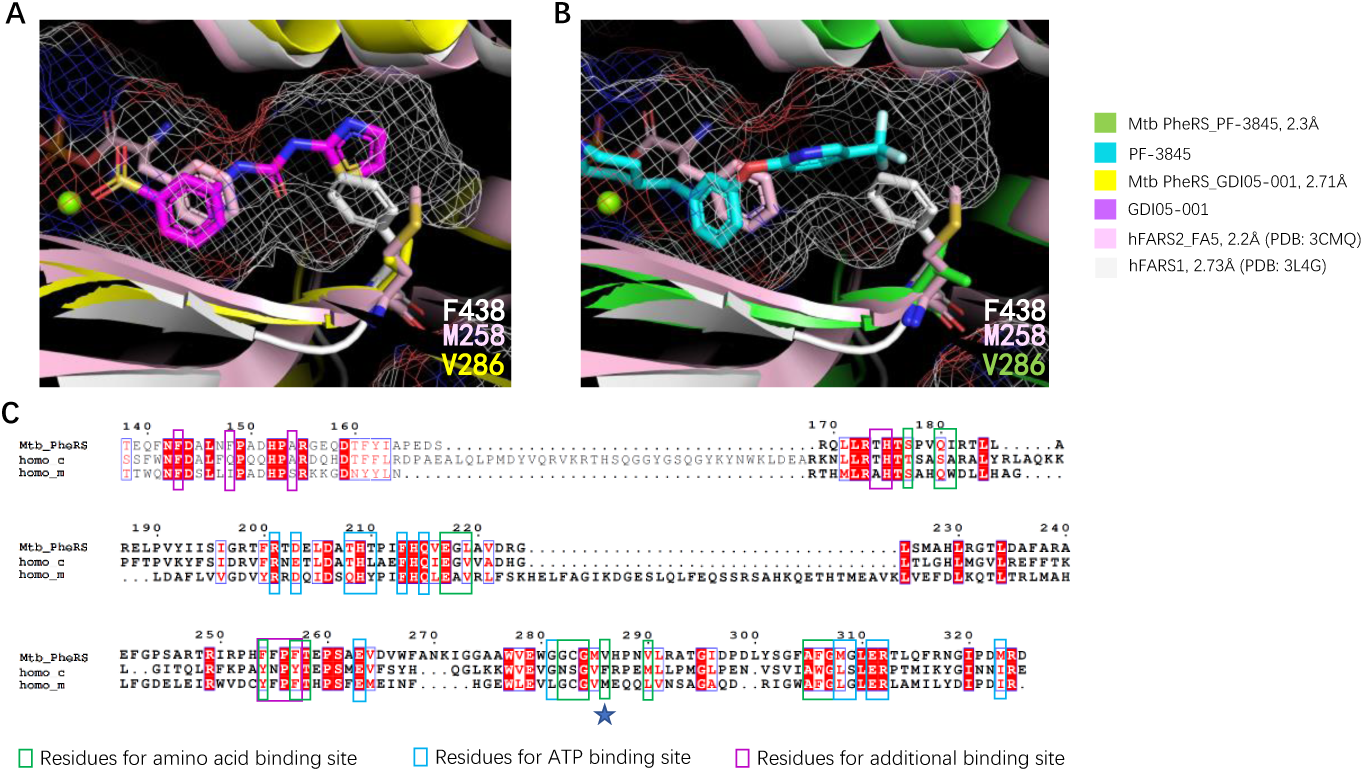
The major difference between the amino acid pockets of Mtb and human PheRSs. A, structure superposition of the amino acid pockets of MtbPheRS-GDI05-001, hFARS1 apo form and hFARS2-Phe-AMP. V286 in Mtb PheRS, F438 in hFARS1 and M258 in hFARS2 are shown in stick. The amino acid pocket of MtbPheRS-GDI05-001 are indicated by mesh. B, structure superposition of the amino acid pockets of MtbPheRS-PF-3845, hFARS1 apo form and hFARS2-Phe-AMP. V286 in Mtb PheRS, F438 in hFARS1 and M258 in hFARS2 are shown in stick. The amino acid pocket of MtbPheRS-PF-3845 are indicated by mesh. C, sequence alignment between Mtb and human PheRS. Key residues forming the three pockets are indicated by boxes in different colors. The most important residue that contributes to the difference of amino acid pocket between Mtb and human is indicated by blue star.

### Interpretation of GDI05-001 NMR and mass spectrum

The ^1^H spectra were recorded in MeOD on Bruker-400 NMR spectrometer. Mass spectrum was collected on Waters UPLC with QDa detector.

^1^H NMR (400 MHz, MeOD) d ppm 2.88 (t, J=7.2 Hz, 2H), 3.20 (t, J=7.6 Hz, 2H), 6.94–7.05 (m, 4H), 7.36 (d, J=3.6 Hz, 1 H), 7.39–7.48 (m, 4H), 7.64 (d, J=3.6 Hz, 1 H), 8.06 (s, 1 H); MS (ESI) m/z calculated for C20H19N5O3S2 [M + H]^+^ 442.1, found 442.3.

## Reference^1^

1. Fischbach, M. A., and Walsh, C. T. (2009) Antibiotics for Emerging Pathogens. Science. 325, 1089–1093

2. Schimmel, P. R., and Söll, D. (1979) Aminoacyl-tRNA synthetases: general features and recognition of transfer RNAs. Annu. Rev. Biochem. 48, 601–648

3. Francklyn, C. S., and Mullen, P. (2019) Progress and challenges in aminoacyl-tRNA synthetase-based therapeutics. J. Biol. Chem. 294, 5365–5385

4. Kwon, N. H., Fox, P. L., and Kim, S. (2019) Aminoacyl-tRNA synthetases as therapeutic targets. Nat. Rev. Drug Discov. 18, 629–650

5. 91. WHO (2020) Global tuberculosis report 2019. Tuberculosis. [online] https://www.who.int/news-room/fact-sheets/detail/tuberculosis (Accessed July 20, 2020)

6. Tenero, D., Derimanov, G., Carlton, A., Tonkyn, J., Davies, M., Cozens, S., Gresham, S., Gaudion, A., Adeep, P., Muliaditan, M., Rullas-Trincado, J., Mendoza-Losana, Alfonso Skingsley, A., and Barros-Aguirre, D. (2019) First-Time-in-Human Study and Prediction of Early Bactericidal Activity for GSK3036656, a Potent Leucyl-tRNA Synthetase Inhibitor for Tuberculosis Treatment. Antimicrob Agents Chemother. 63, 1–15

7. Safro, M., Moor, N., and Lavrik, O. (2005) Phenylalanyl-tRNA Synthetases. in The Aminoacyl-tRNA Synthetases (Ibba, M., Francklyn, C., and Cusack, S. eds), pp. 265–280, Landes Bioscience/Eurekah.com, Georgetown, Texas, USA

8. Dejesus, M. A., Gerrick, E. R., Xu, W., Park, S. W., Long, J. E., Boutte, C. C., Rubin, E. J., Schnappinger, D., Ehrt, S., Fortune, S. M., Sassetti, C. M., and Ioerger, T. R. (2017) Comprehensive essentiality analysis of the Mycobacterium tuberculosis genome via saturating transposon mutagenesis. MBio. 8, 1–17

9. Roy, H., Ling, J., Irnov, M., and Ibba, M. (2004) Post-transfer editing in vitro and in vivo by the β subunit of phenylalanyl-tRNA synthetase. EMBO J. 23, 4639–4648

10. Lechler, A., and Kreutzer, R. (1998) The phenylalanyl-tRNA synthetase specifically binds DNA. J. Mol. Biol. 278, 897–901

11. Gallant, P., Finn, J., Keith, D., and Wendler, P. (2000) The identification of quality antibacterial drug discovery targets: A case study with aminoacyl-tRNA synthetases. Expert Opin. Ther. Targets. 4, 1–9

12. Beyer, D., Kroll, H. P., Endermann, R., Schiffer, G., Siegel, S., Bauser, M., Pohlmann, J., Brands, M., Ziegelbauer, K., Haebich, D., Eymann, C., and Brötz -Oesterhelt, H. (2004) New Class of Bacterial Phenylalanyl-tRNA Synthetase Inhibitors with High Potency and Broad-Spectrum Activity. Antimicrob. Agents Chemother. 48, 525–532

13. Payne, D. J., Gwynn, M. N., Holmes, D. J., and Pompliano, D. L. (2007) Drugs for bad bugs: Confronting the challenges of antibacterial discovery. Nat. Rev. Drug Discov. 6, 29–40

14. Montgomery, J. I., Toogood, P. L., Hutchings, K. M., Liu, J., Narasimhan, L., Braden, T., Dermyer, M. R., Kulynych, A. D., Smith, Y. D., Warmus, J. S., and Taylor, C. (2009) Discovery and SAR of benzyl phenyl ethers as inhibitors of bacterial phenylalanyl-tRNA synthetase. Bioorganic Med. Chem. Lett. 19, 665–669

15. Zhang, Z.-M., Sun, Y., Zhao, L.-L., Wang, L.-N., Wei, Y.-Z., Su, J., Zhang, Y.-Q., and Yu, L.-Y. (2012) The establishment and application of a high throughput screening assay for inhibitors of Mycobacterium tuberculosis phenylalanyl-tRNA synthetase. Microbiol. China. 39, 1437–1446

16. Abibi, A., Ferguson, A. D., Fleming, P. R., Gao, N., Hajec, L. I., Hu, J., Laganas, V. A., McKinney, D. C., McLeod, S. M., Prince, D. B., Shapiro, A. B., and Buurman, E. T. (2014) The role of a novel auxiliary pocket in bacterial phenylalanyl-tRNA synthetase druggability. J. Biol. Chem. 289, 21651–21662

17. Tommasi, R., Brown, D. G., Walkup, G. K., Manchester, J. I., and Miller, A. A. (2015) ESKAPEing the labyrinth of antibacterial discovery. Nat. Rev. Drug Discov. 14, 529–542

18. Hu, Y., Keniry, M., Palmer, S. O., and Bullard, J. M. (2016) Discovery and analysis of natural-product compounds inhibiting protein synthesis in Pseudomonas aeruginosa. Antimicrob. Agents Chemother. 60, 4820–4829

19. Wang, L.-N., Di, W.-J., Zhang, Z.-M., Zhao, L., Zhang, T., Deng, Y.-R., and Yu, L.-Y. (2016) Small-molecule inhibitors of the tuberculosis target, phenylalanyl-tRNA synthetase from Penicillium griseofulvum CPCC-400528. Cogent Chem. 2, 1–9

20. Cowell, A. N., and Winzeler, E. A. (2019) Advances in omics-based methods to identify novel targets for malaria and other parasitic protozoan infections. Genome Med. 11, 1–17

21. Kato, N., Comer, E., Sakata-Kato, T., Sharma, A., Sharma, M., Maetani, M., Bastien, J., Brancucci, N. M., Bittker, J. A., Corey, V., Clarke, D., Derbyshire, E. R., Dornan, G. L., Duffy, S., Eckley, S., Itoe, M. A., Koolen, K. M. J., Lewis, T. A., Lui, P. S., Lukens, A. K., Lund, E., March, S., Meibalan, E., Meier, B. C., McPhail, J. A., Mitasev, B., Moss, E. L.,Sayes, M., Van Gessel, Y., Wawer, M. J., Yoshinaga, T., Zeeman, A. M., Avery, V. M., Bhatia, S. N., Burke, J. E., Catteruccia, F., Clardy, J. C., Clemons, P. A., Dechering, K. J., Duvall, J. R., Foley, M. A., Gusovsky, F., Kocken, C. H. M., Marti, M., Morningstar, M. L., Munoz, B., Neafsey, D. E., Sharma, A., Winzeler, E. A., Wirth, D. F., Scherer, C. A., and Schreiber, S. L. (2016) Diversity-oriented synthesis yields novel multistage antimalarial inhibitors. Nature. 538, 344–349

22. Janes, J., Young, M. E., Chen, E., Rogers, N. H., Burgstaller-Muehlbacher, S., Hughes, L. D., Love, M. S., Hull, M. V., Kuhen, K. L., Woods, A. K., Joseph, S. B., Petrassi, H. M., McNamara, C. W., Tremblay, M. S., Su, A. I., Schultz, P. G., and Chatterjee, A. K. (2018) The ReFRAME library as a comprehensive drug repurposing library and its application to the treatment of cryptosporidiosis. Proc. Natl. Acad. Sci. U. S. A. 115, 10750–10755

23. Martin, F., Eriani, G., Eiler, S., Moras, D., Dirheimer, G., and Gangloff, J. (1993) Overproduction and purification of native and queuine-lacking Escherichia coli tRNAAsp: Role of the wobble base in tRNAAsp acylation. J. Mol. Biol. 234, 965–974

24. Du, X., and Wang, E. D. (2002) Discrimination of tRNALeu isoacceptors by the mutants of Escherichia coli Leucyl-tRNA synthetase in editing. Biochemistry. 41, 10623–10628

25. Ibba, M., Kast, P., and Hennecke, H. (1994) Substrate Specificity Is Determined by Amino Acid Binding Pocket Size in Escherichia coli Phenylalanyl-tRNA Synthetase. Biochemistry. 33, 7107–7112

26. Moor, N., Klipcan, L., and Safro, M. G. (2011) Bacterial and eukaryotic phenylalanyl-tRNA synthetases catalyze misaminoacylation of tRNA Phe with 3,4-dihydroxy-L-phenylalanine. Chem. Biol. 18, 1221–1229

27. Lloyd, A. J., Thomann, H. ulrich, Ibba, M., and Söll, D. (1995) A broadly applicable continuous spectrophotometric assay for measuring aminoacyl-tRNA synthetase activity. Nucleic Acids Res. 23, 2886–2892

28. Baragaña, B., Forte, B., Choi, R., Hewitt, S. N., Bueren-Calabuig, J. A., Pisco, J. P., Peet, C., Dranow, D. M., Robinson, D. A., Jansen, C., Norcross, N. R., Vinayak, S., Anderson, M., Brooks, C. F., Cooper, C. A., Damerow, S., Delves, M., Dowers, K., Duffy, J., Edwards, T. E., Hallyburton, I., Horst, B. G., Hulverson, M. A., Ferguson, L., Jiménez-Díaz, M. B., Jumani, R. S., Lorimer, D. D., Love, M. S., Maher, S., Matthews, H., McNamara, C. W., Miller, P., O’Neill, S., Ojo, K. K., Osuna-Cabello, M., Pinto, E., Post, J., Riley, J., Rottmann, M., Sanz, L. M., Scullion, P., Sharma, A., Shepherd, S. M., Shishikura, Y., Simeons, F. R. C., Stebbins, E. E., Stojanovski, L., Straschil, U., Tamaki, F. K., Tamjar, J., Torrie, L. S., Vantaux, A., Witkowski, B., Wittlin, S., Yogavel, M., Zuccotto, F., Angulo-Barturen, I., Sinden, R., Baum, J., Gamo, F. J., Mäser, P., Kyle, D. E., Winzeler, E. A., Myler, P. J., Wyatt, P. G., Floyd, D., Matthews, D., Sharma, A., Striepen, B., Huston, C. D., Gray, D. W., Fairlamb, A. H., Pisliakov, A. V., Walpole, C., Read, K. D., Van Voorhis, W. C., and Gilbert, I. H. (2019) Lysyl-tRNA synthetase as a drug target in malaria and cryptosporidiosis. Proc. Natl. Acad. Sci. U. S. A. 116, 7015–7020

29. Ahn, K., Johnson, D. S., Mileni, M., Beidler, D., Long, J. Z., McKinney, M. K., Weerapana, E., Sadagopan, N., Liimatta, M., Smith, S. E., Lazerwith, S., Stiff, C., Kamtekar, S., Bhattacharya, K., Zhang, Y., Swaney, S., Van Becelaere, K., Stevens, R. C., and Cravatt, B. F. (2009) Discovery and Characterization of a Highly Selective FAAH Inhibitor that Reduces Inflammatory Pain. Chem. Biol. 16, 411–420

30. Cestari, I., and Stuart, K. (2013) A spectrophotometric assay for quantitative measurement of aminoacyl-tRNA synthetase activity. J. Biomol. Screen. 18, 490–497

31. Moor, N., Kotik-Kogan, O., Tworowski, D., Sukhanova, M., and Safro, M. (2006) The crystal structure of the ternary complex of phenylalanyl-tRNA synthetase with tRNAPhe and a phenylalanyl-adenylate analogue reveals a conformational switch of the CCA end. Biochemistry. 45, 10572–10583

32. Klipcan, L., Levin, I., Kessler, N., Moor, N., Finarov, I., and Safro, M. (2008) The tRNA-Induced Conformational Activation of Human Mitochondrial Phenylalanyl-tRNA Synthetase. Structure. 16, 1095–1104

33. Itoh, Y., Sekine, S. ichi, Kuroishi, C., Terada, T., Shirouzu, M., Kuramitsu, S., and Yokoyama, S. (2008) Crystallographic and mutational studies of seryl-tRNA synthetase from the archaeon Pyrococcus horikoshii. RNA Biol. 5, 169–177

34. Zhou, M., Dong, X., Shen, N., Zhong, C., and Ding, J. (2010) Crystal structures of Saccharomyces cerevisiae tryptophanyl-tRNA synthetase: New insights into the mechanism of tryptophan activation and implications for anti-fungal drug design. Nucleic Acids Res. 38, 3399–3413

35. Collins, L. A., and Franzblau, S. G. (1997) Microplate Alamar blue assay versus BACTEC 460 system for high-throughput screening of compounds against Mycobacterium tuberculosis and Mycobacterium avium. Antimicrob. Agents Chemother. 41, 1004–1009

36. Ahn, K., Smith, S. E., Liimatta, M. B., Beidler, D., Sadagopan, N., Dudley, D. T., Young, T., Wren, P., Zhang, Y., Swaney, S., Van Becelaere, K., Blankman, J. L., Nomura, D. K., Bhattachar, S. N., Stiff, C., Nomanbhoy, T. K., Weerapana, E., Johnson, D. S., and Cravatt, B. F. (2011) Mechanistic and pharmacological characterization of PF-04457845: A highly potent and selective fatty acid amide hydrolase inhibitor that reduces inflammatory and noninflammatory pain. J. Pharmacol. Exp. Ther. 338, 114–124

37. Reshetnikova, L., Moor, N., Lavrik, O., and Vassylyev, D. G. (1999) Crystal structures of Phenylalanyl-tRNA synthetase complexed with phenylalanine and a phenylalanyl-adenylate analogue. J. Mol. Biol. 287, 555–568

38. Mermershtain, I., Finarov, I., Klipcan, L., Kessler, N., Rozenberg, H., and Safro, M. G. (2011) Idiosyncrasy and identity in the prokaryotic phe-system: Crystal structure of E. coli phenylalanyl-tRNA synthetase complexed with phenylalanine and AMP. Protein Sci. 20, 160–167

39. Hatzios, S. K., and Bertozzi, C. R. (2011) The regulation of sulfur metabolism in mycobacterium tuberculosis. PLoS Pathog. 7, 1–8

40. Finarov, I., Moor, N., Kessler, N., Klipcan, L., and Safro, M. G. (2010) Structure of Human Cytosolic Phenylalanyl-tRNA Synthetase: Evidence for Kingdom-Specific Design of the Active Sites and tRNA Binding Patterns. Structure. 18, 343–353

41. Keith, J. M., Apodaca, R., Xiao, W., Seierstad, M., Pattabiraman, K., Wu, J., Webb, M., Karbarz, M. J., Brown, S., Wilson, S., Scott, B., Tham, C. S., Luo, L., Palmer, J., Wennerholm, M., Chaplan, S., and Breitenbucher, J. G. (2008) Thiadiazolopiperazinyl ureas as inhibitors of fatty acid amide hydrolase. Bioorganic Med. Chem. Lett. 18, 4838–4843

42. Bhuniya, D., Kharul, R. K., Hajare, A., Shaikh, N., Bhosale, S., Balwe, S., Begum, F., De, S., Athavankar, S., Joshi, D., Madgula, V., Joshi, K., Raje, A. A., Meru, A. V., Magdum, A., Mookhtiar, K. A., and Barbhaiya, R. (2019) Discovery and evaluation of novel FAAH inhibitors in neuropathic pain model. Bioorganic Med. Chem. Lett. 29, 238–243

43. Grube, C. D., and Roy, H. (2018) A continuous assay for monitoring the synthetic and proofreading activities of multiple aminoacyl-tRNA synthetases for high-throughput drug discovery. RNA Biol. 15, 659–666

44. Hewitt, S. N., Dranow, D. M., Horst, B. G., Abendroth, J. A., Forte, B., Hallyburton, I., Jansen, C., Baragaña, B., Choi, R., Rivas, K. L., Hulverson, M. A., Dumais, M., Edwards, T. E., Lorimer, D. D., Fairlamb, A. H., Gray, D. W., Read, K. D., Lehane, A. M., Kirk, K., Myler, P. J., Wernimont, A., Walpole, C., Stacy, R., Barrett, L. K., Gilbert, I. H., and Van Voorhis, W. C. (2017) Biochemical and structural characterization of selective allosteric inhibitors of the Plasmodium falciparum drug target, prolyl-tRNA-synthetase. ACS Infect. Dis. 3, 34–44

